# Cell Surface Multimeric Assemblies Regulate Canonical and Noncanonical EphA2 Receptor Tyrosine Kinase Signaling

**DOI:** 10.1101/2021.04.11.439330

**Authors:** Xiaojun Shi, Ryan Lingerak, Cameron J. Herting, Yifan Ge, Soyeon Kim, Paul Toth, Carmelle Cuizon, Ji Zheng, Luke Chao, Khalid Sossey-Alaoui, Matthias Buck, Salendra Singh, Vinay Varadan, Juha Himanen, Dolores Hambardzumyan, Dimitar Nikolov, Adam W. Smith, Bingcheng Wang

## Abstract

The EphA2 receptor tyrosine kinase mediates ligand-induced canonical signaling associated with tumor suppression and ligand-independent noncanonical signaling implicated in tumor progression. Using time-resolved fluorescence spectroscopy in live cells, we find that unliganded EphA2 receptors pre-assemble into multimers, which is mediated by two symmetric and one asymmetric interfaces in the ectodomain. Upon ligand binding, EphA2 receptors are further assemble into large clusters that also requires the three interfaces. Functionally, disrupting either the symmetric or asymmetric contacts individually blocks the autorecycling of the EphA2 apo receptor. However, only symmetric contact disruption promotes noncanonical signaling and inhibits ligand-induced catalytic activation and endocytosis, which are associated with increased cell migration *in vitro* and reduced survival in a syngeneic murine glioblastoma model. Our results reveal the pivotal role of EphA2 assembly in dictating canonical vs. noncanonical signaling, and identify the precise molecular interfaces that mediate the formation of the EphA2 signaling clusters.

## Introduction

First cloned from an erythropoietin-producing hepatoma cell line (Hirai et al., 1987), the 16 members of the vertebrate Eph receptors (14 in mammals) constitute the largest family of receptor tyrosine kinases (RTKs). Together with their nine membrane-tethered ephrin ligands, Eph/ephrin form signaling complexes at cell-cell contact sites and regulate a wide range of biological processes during embryonic development and adult physiology (Kania and Klein, 2016) (Nievergall et al., 2012; Pasquale, 2008; Wilkinson, 2001). Aberration of the Eph/ephrin system leads to diverse pathologies, including cancer. EphA2 is the most affected Eph receptor in human malignancies. It is overexpressed in a variety of human solid tumor types, including colon, breast, prostate, and lung cancer, as well as glioblastoma and melanoma, which is often correlated with poor prognosis (Al-Ejeh et al., 2014; Ireton and Chen, 2005; Miao and Wang, 2009; Pasquale, 2010; Wykosky and Debinski, 2008).

Cellular, biochemical and genetic studies have led to the characterization of EphA2 as both a tumor suppressor and an oncogene. This dual function is generally dictated by the ligand binding status. In the presence of ligand, the EphA2 holo receptor, through canonical tyrosine kinase signaling, inhibits the Ras/ERK and PI3K/Akt pathways and inactivates integrin-mediated cell adhesion (Deroanne et al., 2003; Macrae et al., 2005; Miao et al., 2000; Miao et al., 2001; Yang et al., 2011). These properties distinguish EphA2 from prototypic RTKs that activate Akt and ERK upon ligand binding. In fact, activated EphA2 can effectively counter ERK and Akt activation by the EGFR, PDGFR and c-MET receptors upon stimulation with their respective ligands (Miao et al., 2009; Miao et al., 2001; Stallaert et al., 2018). Genetically, *EphA2* deletion in mice dramatically increased susceptibility to carcinogenesis in the mouse skin, supporting an intrinsic tumor suppressive function (Guo et al., 2006).

In its unliganded state, on the other hand, the EphA2 apo receptor becomes a substrate for some serine/threonine kinases, including Akt, p90RSK and PKA, that phosphorylate EphA2 on serine 897 (pS897) (Barquilla et al., 2016; Hamaoka et al., 2016; Miao et al., 2009). Remarkably, this non-canonical signaling event turns EphA2 from a tumor suppressor into an oncogenic protein (Miao et al., 2009). Indeed, recent studies have established S897 phosphorylation as an important regulator of many malignant behaviors, including infiltrative invasion of glioma *in vivo* (Miao et al., 2015), metastases of non-small cell lung cancer (Volz et al., 2020), resistance to BRAF-targeted therapy of melanoma (Azimi et al., 2017) (Paraiso et al., 2015), and chemotherapy resistance of ovarian cancer (Moyano-Galceran et al., 2020). The S897E mutation that simulates pS897-EphA2 in melanoma cells induces mesenchymal-to-amoeboid transition (MAT) and promotes invasion and metastasis(Zhang et al., 2020). In breast cancer cells, pS897 regulates glutamine metabolism (Youngblood et al., 2016) and functions as a mechanosensor for extracellular matrix rigidity to mediate epithelial-mesenchymal transition (EMT) *in vitro* and invasion and metastasis *in vivo* (Fattet et al., 2020). Analysis of human breast cancer showed that vasculogenic mimicry, a poor-prognosis indicator, is also associated with pS897 (Mitra et al., 2020). Together, the recent flurry of reports established an important role of EphA2 noncanonical signaling in promoting malignant progression.

EphA2 is composed of an extracellular domain (ECD), a transmembrane segment (TM), and an intracellular domain (ICD) consisting of a tyrosine kinase domain and a C-terminus sterile α motif (SAM). The ECD includes a ligand-binding domain (LBD), a cysteine-rich domain (CRD) consisting of Sushi and EGF-like motifs, and two fibronectin type III repeats (FN1, FN2) (Himanen and Nikolov, 2003). X-ray crystallography studies of the EphA2 ECD revealed a rigid overall architecture of the unliganded EphA2, undergoing minimal conformational changes upon ligand binding except for a rotation in the FN1-FN2 hinge (Himanen et al., 2010; Seiradake et al., 2010). Ligand engagement induces homotypic interactions between neighboring EphA2 molecules, proposed to induce the formation of large EphA2 clusters intercalated by ephrins (Himanen et al., 2010; Seiradake et al., 2010). FRET-based assays have been used previously to examine the EphA2 spatial organization on cell surface with or without ligand with differing conclusions. An anisotropy-based Homo-FRET assay suggests that EphA2 receptors are monomeric in live cells membranes (Sabet et al., 2015). In contrast, an intensity-based hetero-FRET assays using hypotonically swollen cells reported that unliganded EphA2 formed dimers (Singh et al., 2015).

Here we systematically investigate the molecular assemblies of EphA2 receptors in the cell surface membranes of live cells using a time-resolved fluorescence spectroscopy called pulsed interleaved excitation - fluorescence cross-correlation spectroscopy (PIE-FCCS, **Fig. 1A**) (Bacia and Schwille, 2007; Muller et al., 2005). Our results suggest a new paradigm for RTK assembly into function-defining multimers and sheds light on the molecular basis underlying the dual functions of EphA2 in oncogenesis.

**Figure 1.**
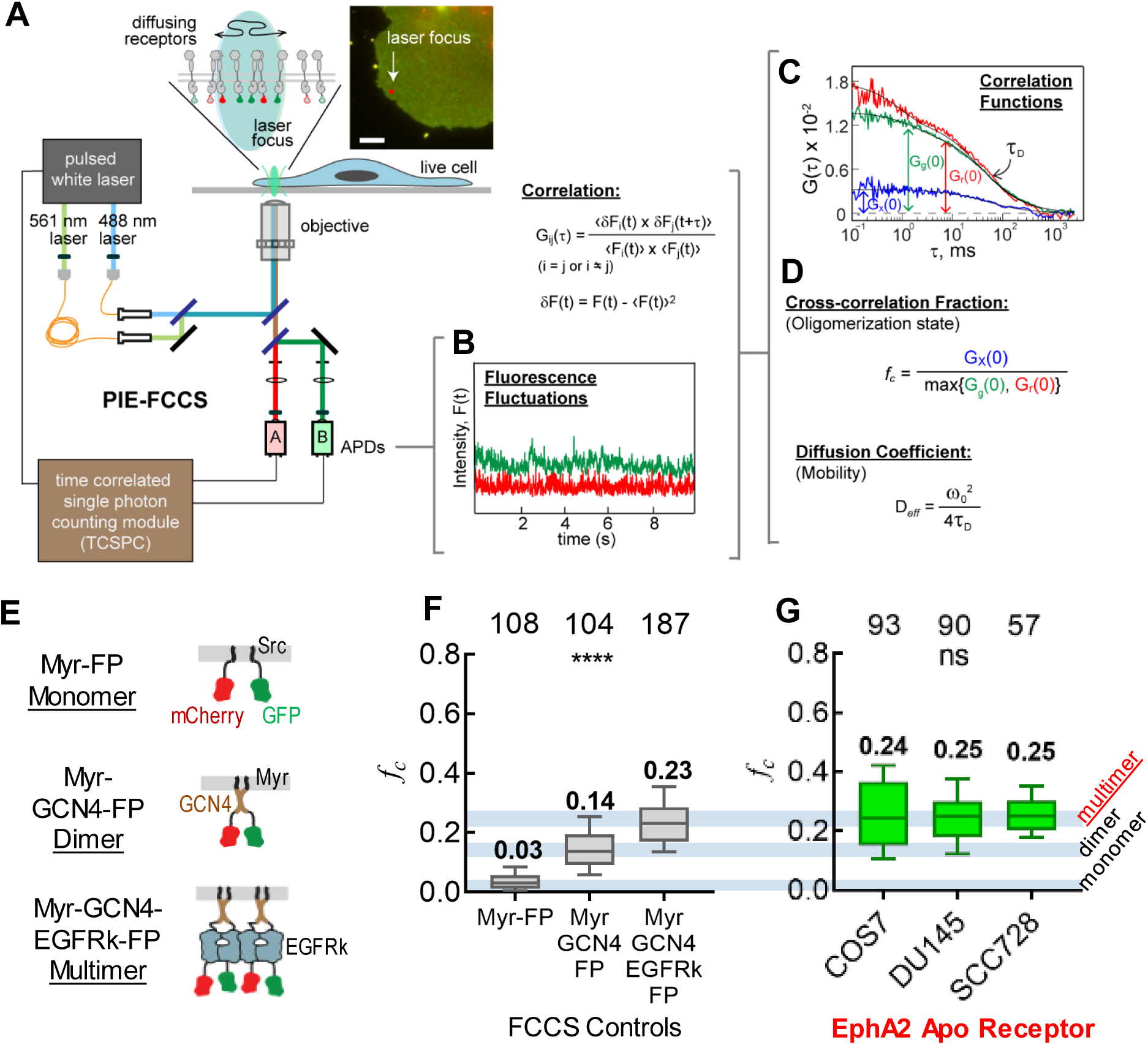
Multimeric Pre-assembly of Unliganded (Apo) EphA2 Detected by PIE-FCCS Measurements. **A)** Schematic of PIE-FCCS instrumentation. Two pulsed lasers are focused on the peripheral membrane of living cells (insert) co-expressing a protein tagged with GFP or mCherry. As the receptors disuse into the focus area, the two laser beams excite the fluorescent probes. Single emitted photons are collected by two APDs and binned into fluorescence intensity signals. The excitation lasers and emitted photons are all synchronized with a TCSPC module. **B)** Fluorescence fluctuation signals collected by APDs after time-gating and binning. **C)** Auto- and cross-correlation functions of the fluorescence intensity data. The green and red lines are the auto-correlation functions corresponding to the GFP- and mCherry-labeled receptors respectively. The blue line is the cross-correlation function. **D)** The relative cross-correlation value (*f_c_*) is calculated from the correlation function amplitudes at τ = 0, (G(0)), and is related to the oligomerization state of the labelled proteins. The effective diffusion coefficient (*D_eff_*) is calculated from the dwell time (τ_D_) of receptor assemblies within the laser focus, which is related to the mobility of the labeled proteins. Because the oligomerization state of the proteins has direct effect on their mobility, reporting both parameters provides a more complete view of membrane protein oligomerization in live cells. **E)** Diagram of oligomerization control constructs. The monomeric control, Myr-FP, is a co-expression of GFP and mCherry, each fused separately to c-Src membrane localization sequence. The dimeric control, Myr-GCN4-FP, has the leucine zipper dimerization motif of GCN4 fused to GFP/mCherry and the c-Src membrane localization sequence. The multimeric control, Myr-GCN4-EGFRk-FP, has the self-dimerizing kinase domain of EGFR introduced after the GCN4 motif. **F)** Single-cell cross-correlation values (*f_c_*) for each of the control constructs taken concurrently with the EphA2 data. **G)** Single-cell cross-correlation values (*f_c_*) of unliganded (Apo) EphA2 in the plasma membranes of three cell lines: COS7, DU145, and SCC728. Each data point is the average of five 10 s measurements performed on one cell. Cross-correlation values are plotted as box-whisker plots. The boxes represent third quartile, median and first quartile. The whiskers indicate 10-90^th^ percentile. The numbers on top of the box plots are the total number of cells used. One-way ANOVA tests are performed to obtain p values (****: p < 0.0001; ns: not significant).

## Results

### 1. Time-Resolved Fluorescence Spectroscopy Reveals Pre-Assembly of EphA2 Apo Receptor into Multimers in Live Cells

To understand the molecular basis of canonical and noncanonical signaling by EphA2, we interrogated the spatiotemporal assemblies of EphA2 in live cells using a dual-color, time-resolved fluorescence spectroscopy called PIE-FCCS (**Methods**). In PIE-FCCS two pulsed lasers are overlapped in space but interleaved in time so that each detected photon can be assigned to an excitation source (**Fig. 1A**) (Muller et al., 2005) (Christie et al., 2020). Data are collected from single cells by focusing the lasers on the plasma membranes (**Fig. 1A, insert**). Photons emitted from labeled receptors diffusing through the laser focus are collected, binned and time-gated to yield fluorescence fluctuation signals (**Fig. 1B**). The signals are then transformed into auto-and cross-correlation functions. From these correlation functions (**Fig. 1C**), the degree of oligomerization and mobility of the receptors can be quantified (**Fig. 1D** and Method) by monitoring the diffusion of receptors and specifically, the collective diffusion of oligomerized receptors. The diffusion-based approach of PIE-FCCS is less affected by dynamic collisions, crowding and complex photophysics that usually complicate intensity-based fluorescence assays. Hence, PIE-FCCS can provide a rigorous readout of receptor oligomerization in live cell membranes. Control systems consisting of membrane-bound protein monomers, dimers and multimers (**Fig. 1E, F and S1A**) were measured concurrently with EphA2 samples to provide a cross-correlation value (*f_c_*) scale for quantification of the oligomerization state of receptors.

To determine the oligomerization state of EphA2, we co-expressed EphA2 tagged with GFP or mCherry in COS7 cells, a model cell line used in previous PIE-FCCS studies (Endres et al., 2013). Cells expressing EphA2 ranging from 200 to 2000 receptors/μm^2^ (**Fig. S1C**) were chosen, which is compatible with the physiological level of EphA2. Unexpectedly, in the absence of ligand binding, the *f_c_* values for the EphA2 apo receptor **(Fig. 1G**) were significantly larger than those for the dimer control and were more comparable to those for the multimer control (**Fig. 1F**). Similar results were obtained in DU145 human prostate cancer cells. These observations suggest that unliganded EphA2 apo receptors self-assembled predominantly into multimers.

EphA2 is widely expressed in many epithelial cell types (Lindberg and Hunter, 1990). To control for potential interference from endogenous EphA2 and the closely related EphA1, we used a mouse cutaneous squamous cell carcinoma cell line (SCC728) established from an *EphA2* and *EphA1* double knockout mouse (**Methods**). PIE-FCCS measurements show that the *f_c_* values in SCC728 cells are at the same levels as those in COS7 and DU145 cells, consistent with multimerization of EphA2 apo receptor. To our knowledge, this is the first multimeric assembly observed for an RTK in its ligand-free state. By contrast, the unliganded EGFR in the same PIE-FCCS assay is predominantly monomeric(Endres et al., 2013).

### 2. Three Distinct Interfaces in the Ectodomain Mediate the Multimeric Assembly of EphA2

#### Two Symmetric Head-to-Head (HH) Interfaces, LBD-LBD and Sushi-Sushi, are Required for Apo EphA2 Multimeric Assembly

To elucidate the molecular basis of apo EphA2 multimerization, we combined PIE-FCCS with mutagenesis based on interfaces previously observed in the EphA2 crystal structures(Himanen et al., 2010; Seiradake et al., 2010). We reasoned that these interactions could participate in the assembly of the EphA2 apo receptor in physiological settings. We first investigated the symmetric head-to-head (HH) Eph-Eph contact site located at the C-terminus of the ligand-binding domain (LBD). This interface is formed by hydrogen bonds and Van der Waals interactions centered around residues D129 and G131, and buries a total molecular surface area of ∼1100 Å^2^ (**Fig. 2A**) (Himanen et al., 2010; Seiradake et al., 2010). We mutated both residues, D129N and G131S, and denoted this mutation as ‘LBD’ (**Fig. S2A**). PIE-FCCS measurements showed that the *f_c_* values for the LBD mutation were significantly reduced relative to those for the WT receptor (**Fig. 2B left**). This observation suggests that disruption of the LBD-LBD interface reduces the degree of EphA2 multimerization, which is supported by an increase in receptor mobility (**Fig. 2B right**) that can be ascribed to decreased friction within the membrane for smaller receptor assemblies relative to larger ones. This is evidence that the LBD interface participates in the multimerization of the EphA2 apo receptor on the cell surface.

**Figure 2.**
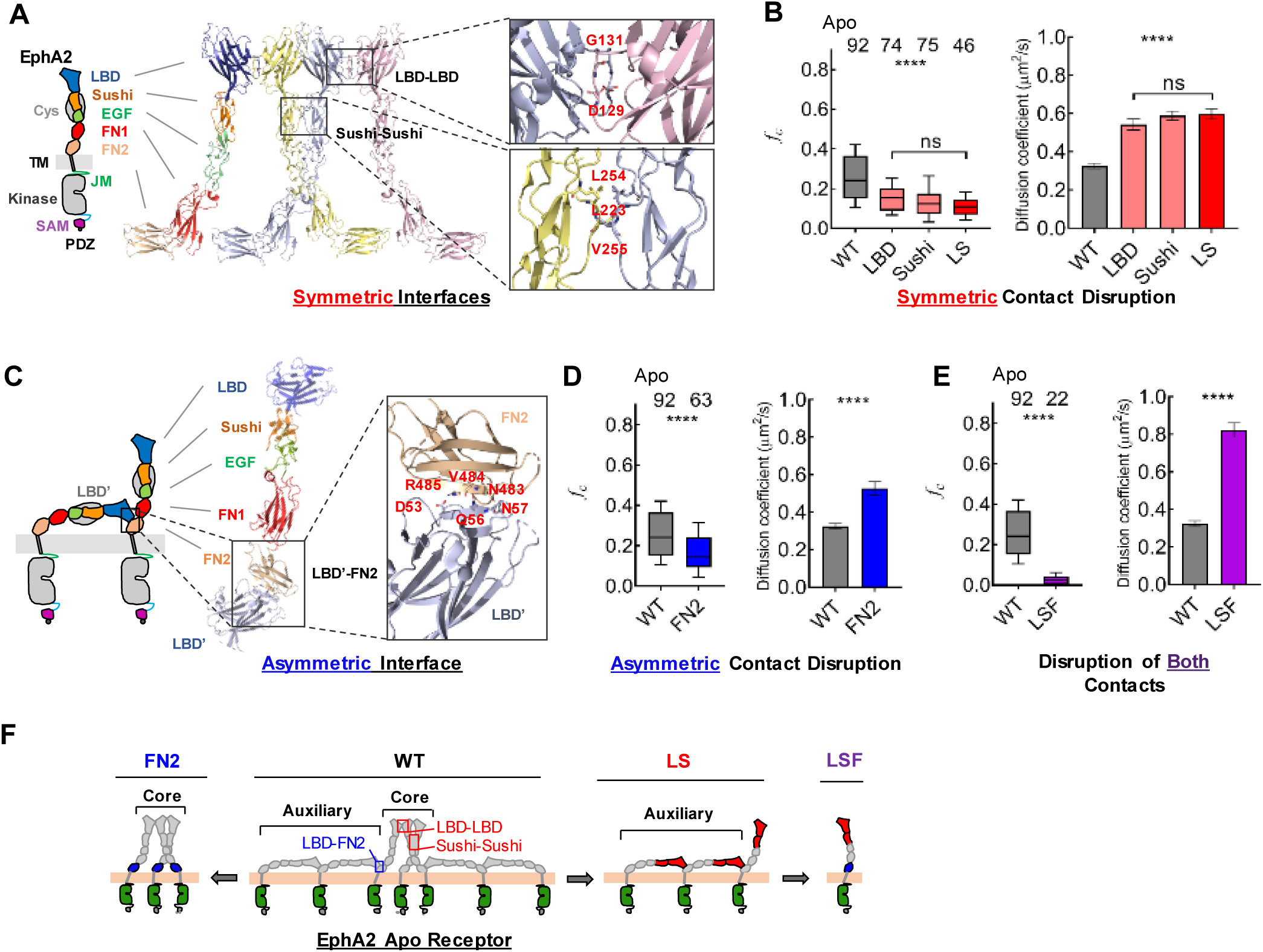
Multimerization of Apo EphA2 is Mediated by Symmetric Head-Head and Asymmetric Head-Tail Interfaces. **A)** EphA2 domain structure. LBD, ligand-binding domain; Cys, cysteine-rich domain; Sushi; EGF, epidermal growth factor-like domain; FN, fibronectin domain; TM, transmembrane region; JM, juxtamembrane segment; SAM, sterile α motif; PDZ, Psd-95, Dlg and ZO1 domain-binding motif. Crystal structure of EphA2 ectodomain highlighting LDB-LBD and Sushi-Sushi symmetric interfaces. Inserts: Detailed view of LDB-LBD and Sushi-Sushi interfaces. Residues that mediate interactions are labeled. **B)** *left:* Cross-correlation values of WT and mutated EphA2 apo receptors. WT, wild-type EphA2; LBD, LBD-LBD interface impaired EphA2; Sushi, Sushi-Sushi interface impaired EphA2; LS, both Sushi-Sushi and LBD-LBD interfaces impaired EphA2. *right:* Diffusion coefficients of apo EphA2 receptors with disruptions at symmetric contacts. The error bars represent SEM from the number of cells analyzed as indicated on the top in the left panel. **C)** *Left:* Model of two apo EphA2 molecules adapting asymmetric contact through FN2 and LBD. *Right*: Crystal structure of asymmetric contact. *Insert*: Detailed view of LBD-FN2 interface. Residues that mediate interactions are labeled. **D)** Cross-correlation (left) values and diffusion coefficients (right) of WT vs. FN2 mutant apo EphA2 with disruption of the asymmetric LBD-FN2 contact. **E)** Cross-correlation values (left) and diffusion coefficients of WT vs. LSF mutant apo EphA2 with disruptions at both symmetric and asymmetric contacts. The cross-correlation values of LSF are close to zero, indicating LSF is mostly monomeric. **F)** Schematic illustration of the molecular assemblies of wild-type EphA2 and mutants at apo state. The one-way ANOVA tests were performed to obtain p values (****: p < 0.0001; ns: not significant).

Next, we examined the leucine-zipper-like symmetric interface involving the Sushi domains of adjacent receptors, which is composed of hydrophobic contacts burying a surface area of ∼850 Å^2^ (**Fig. 2A**) (Himanen et al., 2010; Seiradake et al., 2010). L223, L254 and V255 were mutated to arginine to disrupt this Sushi-Sushi interface (denoted ‘Sushi’, **Fig. S2A**). The median *f_c_* values decreased from 0.24 to 0.12 (**Fig. 2B left**), suggesting a reduction in the degree of multimerization, which is supported by an increase in mobility (**Fig. 2B right**). Thus, similar to the LBD interface, the Sushi interface contributes to the multimerization of the EphA2 apo receptor on the cell surface.

Finally, we evaluated the effects of simultaneously disrupting both the LBD and Sushi interfaces. The *f_c_* values for the compound mutation, named ‘LS’ (**Fig. S2A**), were similar to the LBD or Sushi mutations alone (**Fig. 2B left**), suggesting that disruption of either symmetric interface alone was sufficient to destabilize the symmetric contacts in the apo EphA2 receptors. However, LS still has *f_c_* values greater than the zero value of the monomeric controls (0.11 vs. 0.03, **Fig. 2B left** and **Fig. 1F**), which suggests that additional interface(s) contribute to the multimeric assembly of Apo EphA2.

In searching for other potential ectodomain interactions, we investigated a third putative symmetric interface observed in one of the EphA2/ephrin-A5 crystal structures, which is located in the first type III FN domain (FN1) (Seiradake et al., 2010). We disrupted this FN1-FN1 interface by generating a P378E mutation named ‘FN1’ (**Fig. S2A**) (Seiradake ^e^_t al., 201_^0^). Interestingly, no changes of the *f_c_* values were observed in PIE-FCCS measurements of FN1 (**Fig. S2B**). The mobility of FN1 was also similar to this of WT EphA2 (**Fig. S2C**). We conclude that the FN1-FN1 interaction is unlikely to play a significant role in the formation of unliganded multimers of EphA2.

#### A Novel LBD-FN2 Interface Mediates Head-Tail Asymmetric Assembly of EphA2

It was previously suggested (Nikolov et al., 2014), based on the structures of the unliganded EphA2 (Himanen et al., 2010) and EphA4 (Xu et al., 2013), that a novel Eph-Eph interface formed between the LBD and the second FNIII domain (FN2) (**Fig. 2C**) of adjacent apo receptors *in cis* (on the same cell), might contribute to Eph signaling initiation. This LBD-FN2 interface (**Fig. 2C**) in EphA2 (∼980 Å^2^ total buried surface area) involves several salt bridges and hydrogen bonds, including a salt bridge between D53 and R485, hydrogen bonding of the side chain of Q56 with the main chain of V484, as well as the main chain of N57 with the side chain of N483 on the neighboring LBD and FN2 domains respectively (**Fig. 2C**). To determine the contribution of this novel interface to EphA2 apo receptor multimerization, a double N483L/R485E mutation was made to disrupt the LBD-FN2 interactions (denoted ‘FN2’, **Fig. S2A**). This mutation led to a decrease in the *f_c_* values from 0.24 to 0.15 for EphA2 apo receptors (**Fig. 2D left**), while increasing the mobility of the receptors (**Fig. 2D right**). Thus, the asymmetric LBD-FN2 interaction also contributes to the preassembly of apo EphA2 into multimers.

#### Disrupting Both Symmetric and Asymmetric Interfaces Turns the EphA2 Apo Receptor Multimers into Monomers

We next simultaneously disrupted all three interfaces (denoted as ‘LSF’, **Fig. S2A**). As shown in **Fig. 2E**, the *f_c_* values for the LSF mutation are diminished close to zero, similar to that of the monomeric controls (**Fig. 1F**), accompanied by dramatically increased mobility (**Fig. 2E right**) (Note the color coding of the mutations in Figures: **LBD**, **Sushi** and **LS** in two shades of red; **FN** in blue, and **LSF** in purple). This demonstrates that the LSF EphA2 mutant exists as a monomer. While additional Eph-Eph interfaces with relatively minor contributions cannot be ruled out, we conclude that the multimeric assembly of the EphA2 apo receptor is primarily mediated by the symmetric LBD-LBD and Sushi-Sushi contacts, as well as the asymmetric LBD-FN2 interactions.

A schematic consistent with the PIE-FCCS results is presented in **Fig. 2F**. EphA2 apo receptors assemble into multimers through a “core” assembly of Eph molecules connected by symmetric HH interfaces (LBD-LBD, Sushi-Sushi) flanked by “auxiliary” arms formed by asymmetric HT (LBD-FN2) interactions. Disruption of symmetric (LS) and asymmetric (FN2) interfaces leads to smaller oligomerization sustained by auxiliary and core assemblies, respectively.

#### The Intracellular Domains of EphA2 Apo Receptors in the Auxiliary Arms Are Farther Apart from Each Other Than Those in the Core

At 146 Å, the ECD of EphA2 is both long and rigid(Himanen et al., 2010). As such, one prediction based on our model (**Fig. 2F)** is larger distances between the adjacent ICDs of EphA2 in the auxiliary arms compared with those in the core. To test this, we used fluorescence anisotropy measurements to detect fluorescence resonance energy transfer among identical GFPs (homoFRET) resulting from proximity between GFP-tagged cytoplasmic tails. Closer proximity among GFPs leads to stronger homoFRET and lower fluorescence anisotropy(Altman et al., 2007; Hunt et al., 2012; Ross et al., 2018). We found that the anisotropy of FN2 is significantly smaller than that of LS (**Fig. S2E**), despite the similar degree of oligomerization from PIE-FCCS measurements (**Fig. 2B and D, Fig. S2D**). This observation indicates closer proximity between the ICDs in the FN2 mutant than those in the LS mutant, providing further support for the model shown in **Fig. 2F**. By locking EphA2 in monomer, the ICDs in LSF mutant are farthest apart and therefore displayed the highest anisotropy values(**Fig. S2E, F**). Having a mixture of both contacts, WT EphA2 shows a medium anisotropy value (**Fig. S2E**).

### 3. Ephrin-A1 ligand Is Monomeric on Cell Surface

Before investigating how ligand binding affects EphA2 assembly, we used PIE-FCCS to assess how the ephrin-A1 ligand is presented on live cell surfaces. This is important because ephrin-A ligand organization under physiological conditions is yet to be determined, and inconsistent results have been reported as to whether ephrin-A ligands have to be artificially dimerized or multimerized to achieve full EphA2 activation (Davis et al., 1994; Wykosky et al., 2008; Xu et al., 2011). GFP and mCherry tags were inserted after the signal sequence of ephrin-A1 and its oligomerization status was measured by PIE-FCCS. Interestingly, ephrin-A1 displayed an *f_c_* values close to zero with high mobility (**Fig. 3A**), similar to the monomer control (**Fig. 1F**), suggesting that ephrin-A1 exists as a monomer on the surface of live cells. These findings have significant implications in understanding bidirectional signaling and function of Eph-Ephrin system in development and diseases.

**Figure 3.**
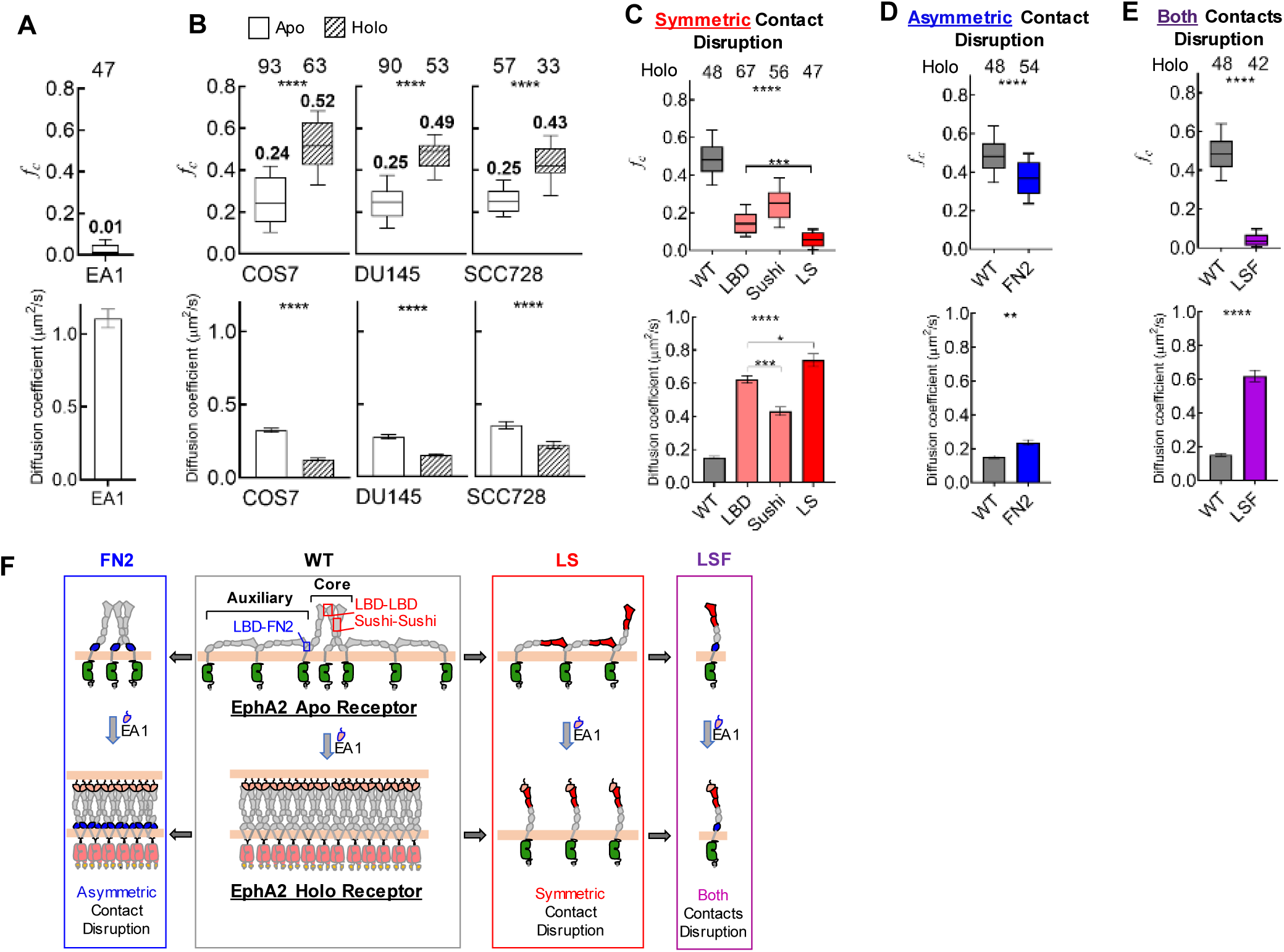
Symmetric and Asymmetric Contacts Contribute to Ligand-induced Clustering of EphA2. **A)** Cross-correlation values (top) and diffusion coefficients (bottom) of ephrinA1 in the plasma membranes (PM) of COS7 cells. **B)** Cross-correlation values (top) and diffusion coefficients (bottom) of unliganded (apo) and dimeric EA1-Fc ligand-stimulated (holo) EphA2 in the three cell types. **C)** Cross-correlation values (top) and diffusion coefficients (bottom) of monomeric ephrinA1 (mEA1) stimulated (holo) EphA2 with disruption at symmetric contacts (LBD, Sushi and LS). **D)** Cross-correlation values (top) and diffusion coefficients (bottom) of holo EphA2 construct (FN2) with disruption of the asymmetric contact. **E)** Cross-correlation values (top) and diffusion coefficients (bottom) of holo EphA2 with disruption at both symmetric and asymmetric contacts. The one-way ANOVA tests were performed to obtain p values (****: p < 0.0001; ***: p < 0.001; **: p < 0.01; *: p < 0.05). **F)** Schematic diagram of the molecular assemblies of wild-type EphA2 and mutants that have symmetric and/or asymmetric contacts disrupted.

#### Both Monomeric and Artificially Dimerized Ephrin-A1-Fc Induce Higher Order EphA2 Receptor Clustering

Based on the results above, both monomeric ephrin-A1 (mEA1) and commonly utilized dimeric ephrin-A1-Fc were examined for effects on EphA2 assembly with PIE-FCCS. Upon stimulation with ephrin-A1-Fc, which is known to cause robust activation of EphA2 (Davis et al., 1994; Miao et al., 2000), we detected dramatically increased *f_c_* values, far above those for the multimeric apo receptor, in all cell lines tested, including COS7, DU145, as well as SCC728 (**Fig. 3B**, top). These increased *f_c_* values, together with a significant decrease in receptor mobility (**Fig. 3B**, bottom), suggest the formation of higher-order clusters of EphA2 receptors. Although the sizes of the clusters cannot be precisely determined by PIE-FCCS, the results are consistent with holo EphA2 undergoing ligand-induced lateral condensation and aggregation into high order clusters also observed in the crystallographic studies (Himanen et al., 2010; Seiradake et al., 2010). Interestingly, treatment with physiologically relevant monomeric mEA1 also induced the large clusters of WT EphA2 to a similar degree as dimeric ephrin-A1-Fc (**Fig. 3C**, WT; **Fig. S3A**). Since we determined that ephrin-A1 is a monomer in live cell membranes (**Fig. 3A**), we suspected that the artificially dimerized ephrin-A1-Fc might force non-physiologically relevant clustering of EphA2 and, therefore, mEA1 is used for the following PIE-FCCS experiments.

#### The Symmetric Interfaces Mediate Ligand-Induced EphA2 Receptor Clustering, While the Asymmetric Interactions Are Outcompeted by Ligand

Next mEA1 was used to stimulate EphA2 containing mutations that disrupt the three multimerization interfaces. PIE-FCCS measurements show that disrupting each of the two symmetric interfaces individually reduces the degree of higher-order cluster formation induced by mEA1, with more pronounced effects in the LBD mutation (**Fig. 3C**). Intriguingly, mEA1 stimulation of the combined LS mutant, which disrupts both symmetric LBD and Sushi interfaces, caused a further reduction of the *f_c_* values from 0.11 for the apo receptor (**Fig. 2B**) to close to zero, in the range expected of a monomer (**Fig. 3C**). However, viewed in the context of the relative affinities of the EphA2 LBD for ephrin-A1 vs. FN2, and the fact that the LBD-FN2 and LBD-ephrin-A1 interfaces overlap (Nikolov et al., 2014) (**Fig. S3B**), the finding was actually expected. As illustrated in **Fig. 3F**, the weaker LBD-FN2 interaction can be outcompeted by the much stronger LBD-mEA1 interaction, leading to the formation of a ligand-receptor complex that moves in unison and thus displays *f_c_* values of a monomer. Indeed, this LBD-FN2 interface is not present in the crystal structures of 1:1 Eph/ephrin complexes (Himanen et al., 2010) (Seiradake et al., 2010).

Stimulation of EphA2 harboring the FN2 mutation, which disrupts the asymmetric interface, showed a significant but milder decrease in the degree of clustering as compared to the WT EphA2 (**Fig. 3D**). It has been suggested that these asymmetric contacts help bring more Eph receptors together prior to cell-cell contact-induced Eph-Ephrin engagement, ensuring fast and efficient ligand-induced clustering and condensation (Nikolov et al., 2014). Thus, the disruption of the asymmetric Eph-Eph contacts would reduce the availability of Eph receptors in the immediate proximity of the forming Eph/ephrin higher-order clusters.

LSF mutant with all three interfaces disrupted stayed as a monomer in the presence of mEA1 (**Fig. 3E**). Of note, none of the mutations affected the ligand binding between EphA2 and ephrin-A1 on the cell surface (**Fig. S3C**), indicating the effects are specifically associated with Eph oligomerization status. Confirming our concerns that the artificially dimerized ligand may force non-physiological receptor dimerization/multimerization, stimulation with dimeric ephrin-A1-Fc caused larger *f_c_* values for both LS and LSF mutants (**Fig. S3D-G**) than those induced by mEA1 (**Fig. 3C-E**).

The ligand-induced changes EphA2 oligomerization are schematically summarized in **Fig. 3F**. Ligand binding to WT EphA2 disrupts the LBD-FN2 interface in the auxiliary flanks. The ensuing conformational changes at the FN1-FN2 hinge leads to lateral clustering of EphA2 holo receptors and recruitment of additional receptors through symmetric interfaces(Seiradake et al., 2010), culminating in compaction of the EphA2 receptors into higher order clusters intercalated with ephrin ligands (**Fig. 3F**, WT). The LS mutant participates in only LBD-FN2 interactions that can be outcompeted by the incoming ligand, resulting in a mostly monomeric EphA2 (**Fig. 3F**, LS). In the FN2 mutant, only Eph core assembly remains intact, permitting robust clustering by the ligand, but to a smaller extent than in the WT EphA2 (**Fig. 3F**, FN2) due to the lack of auxiliary EphA2. Disrupting all three interfaces renders EphA2 in a fully monomeric state (**Fig. 3F**, LSF), irrespective of ligand binding.

**Table S1** summarizes the cross correlation values (fc), diffusion coefficient, GFP lifetime and anisotropy for WT and all four mutant EphA2 (Sushi, LBD, LS and LSF) in the absence (apo) and presence (holo) of monomeric or dimeric ligand.

### 4. EphA2 Receptor Oligomerization Regulates Canonical and Noncanonical Signaling as well as Constitutive Recycling and Endocytosis

#### The Symmetric Interfaces Are Necessary and Sufficient for Ligand-Dependent Canonical Signaling

Ligand stimulation induces Eph canonical signaling characterized by tyrosine phosphorylation on the di-tyrosine motif in the juxtamembrane (JM) segment that is conserved among most Eph receptors(Pasquale, 2005). The SCC728 cells with EphA1/EphA2 DKO provide a clean platform to examine the roles of multimerization in EphA2 canonical signaling. Expression of exogenous WT EphA2 in SCC728 cells showed strong canonical signaling after 15 min ligand stimulation (**Fig. 4A**, WT, pY-EphA/B). Longer treatment caused degradation of EphA2. Similar results were observed in HEK293, GSC827, PC3 and SCC748 cells expressing exogenous WT EphA2 (**Fig. S4A-D**). The FN2 mutant remained similarly responsive (**Fig. 4A**, FN2), suggesting the LBD-FN2 interactions are dispensable for EphA2 canonical signaling. In contrast, perturbations of the two symmetric interfaces alone (LBD or Sushi) or in combination (LS), largely blocked ligand-induced EphA2 tyrosine phosphorylation (**Fig. 4A**). Perturbation of all three interfaces (LSF) caused further reduction in canonical signaling in all cell types tested, most notably in HEK293 cells (**Fig. 4A** and **Fig. S4**). We conclude that the symmetric interfaces are indispensable for ligand-induced canonical signaling.

**Figure 4.**
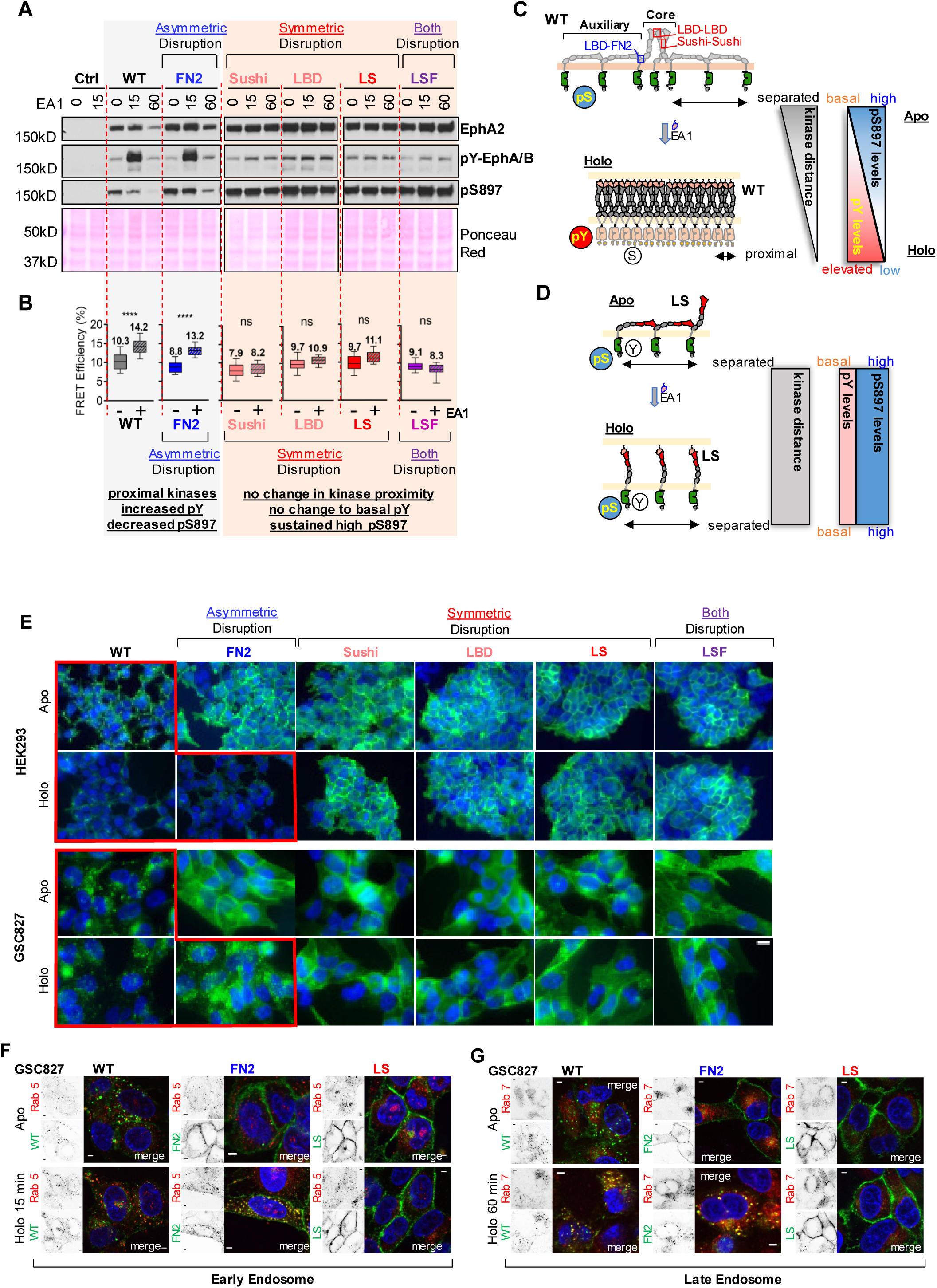
Symmetric and Asymmetric Contacts Modulate Signaling and Endocytosis of EphA2. **A)** WT and the indicated mutant EphA2 are expressed in SCC728 cells. Following stimulation with ephrinA1-Fc, cells are lysed and subjected to immunoblot. Ponceau Red staining is used as loading control. Similar results are obtained from GSC827, HEK293, SCC748, PC3 and 283LM cells (**Fig. S4A-D**) and COS7, DU145 and GSC1816 cells (data not shown). **B)** FRET efficiency of WT and mutant EphA2 before and after ligand stimulation is calculated based on the lifetime of monomer control. The one-way ANOVA tests were performed to obtain p values (****: p < 0.0001). **C, D)** Schematic diagram of signaling and change of kinase proximity of WT **(C)** and LS **(D)** EphA2. **E)** Epi-fluorescence images of HEK293 (top) and GSC827 (bottom) expressing WT and mutant EphA2 at unliganded (apo) and EA1-stimulated (holo) states. **F, G)** Confocal images of GSC827 cells expressing WT or mutant EphA2 stained for Rab 5 **(F)** and Rab 7 **(G)** following 15 and 60 minutes stimulation with eohrin-A1, respectively. The EphA2 constructs are tagged with GFP and the staining of Rab proteins is visualized with Alexa Fluor 563 conjugated secondary antibody. The nuclei are stained with DAPI. Separated images of EphA2 constructs (GFP) and Rab proteins (Alexa Fluor 563) are shown in inverted format. Merged images of the cells are shown in colors. The yellow punctuated features indicating the co-localization of EphA2 and Rab protein. All scale bars are 5 μm.

#### EphA2 Multimerization Through the ECD Contributes to ICD Juxtaposition and Catalytic Activation

Generally, ligand-binding by RTKs induces conformational changes in the ECDs, which are propagated to the ICDs, leading to kinase domain approximation, catalytic activation and tyrosine transphosphorylation (Endres et al., 2014; Schlessinger, 2014). To investigate whether EphA2 multimerization mediated by the three interfaces in the ECD contributes to the ICD rearrangement and catalytic activation upon ligand stimulation, we examined the fluorescence lifetime of GFP recorded during PIE-FCCS measurements. A decrease in GFP fluorescence lifetime suggests increased FRET efficiency between the C-terminus GFP and mCherry tags, indicating an increased proximity between the ICDs. To test this concept, a panel of controls were engineered where GFP and mCherry are kept either close together or farther apart (**Fig. S4E**). The GFP lifetime measurements revealed a close correlation of GFP lifetime with the distances among the engineered proteins (**Fig. S4F**). In cells expressing WT or FN2 mutant, ligand stimulation resulted in increased FRET efficiency (**Fig+. 4B**), coinciding with increased tyrosine phosphorylation (**Fig. 4A**). In contrast, mutations affecting the two symmetric interfaces alone or in combination showed little changes in FRET efficiency upon ligand stimulation (**Fig. 4B**, Sushi, LBD and LS) in keeping with lack of tyrosine phosphorylation. These results indicate that ligand-induced conformational changes in EphA2 ECD indeed propagate to the ICDs through the symmetric interfaces to bring the kinase domain into close proximity to cause catalytic activation (**Fig. 4C**). The lifetime results with ligand-bound holo receptor complement those from the homoFRET measurements of unliganded apo receptors, supporting for an overall model of EphA2 assembly (**Fig. 3F**).

#### The LBD-FN2 Asymmetric Interactions Facilitate Ligand-Independent Non-Canonical Signaling

EphA2 noncanonical signaling is marked by serine 897 phosphorylation (pS897)(Miao et al., 2009). We found that perturbation of either asymmetric or symmetric interfaces increased basal pS897 (**Fig. 4A, Fig. S4A-D**). Consistent with earlier report(Miao et al., 2009); ligand stimulation reduced phosphorylation on S897 of WT EphA2, an effect that is retained in the FN2 mutant. In contrast, pS897 levels remained high and unchanged upon sustained ligand exposure for the symmetric interface mutants (LBD, Sushi and LS). Since the latter mutations disrupt the core assembly while leaving the auxiliary flanks intact (**Fig. 3F**), these results show that the asymmetric LBD-FN2 interactions facilitates noncanonical signaling. This is possibly achieved by keeping ICDs spatially distant in the auxiliary flanks, a notion supported by the homoFRET measurements described above (**Fig. S2E.F**).

#### Effects of the EphA2 Multimeric Preassembly on the Ligand-Independent Constitutive Recycling

Like other RTKs, the EphA2 apo receptor undergoes ligand-independent auto-activation and recycling to ensure proper cell surface expression and optimal response to ligand stimulation (Sabet et al., 2015). This is evident in images of HEK293 cells and GSC827 glioma stem cells expressing WT EphA2. Without exposure to ligand, we observed two populations of WT EphA2 apo receptor, one on the plasma membrane and the other in the cytosol (**Fig. 4E, Fig. S4G**). Remarkably, mutations perturbing any of the three Eph-Eph interfaces individually or in combinations led to profound accumulation of EphA2 on the cytoplasmic membrane and reduction of the cytosolic pool (**Fig. 4E, Fig. S4G**), suggesting that the multimeric preassembly of the EphA2 apo receptor is required for autorecycling. The increased cytoplasmic membrane retention also makes the mutant more accessible to pS897 phosphorylation by Akt, p90RSK and PKA kinases, contributing to the increased basal pS897 (**Fig. 4A**). Upon stimulation with ephrin-A1, WT and FN2 EphA2 became degraded, whereas no degradation was seen with the symmetric interface mutants (**Fig. 4E**), which corroborates well with the immunoblot results (**Fig. 4A**).

#### Effects of Eph Oligomerization on Ligand-Induced Endocytosis

Endocytosis plays an essential role in signaling by RTKs (Miaczynska, 2013), including Eph kinases (Pitulescu and Adams, 2010). To investigate how EphA2 multimerization impacts endocytosis, we examined ligand-induced EphA2 trafficking in GSC827 cells expressing WT EphA2 or with FN2 and LS mutations. In agreement with earlier reports (Boissier et al., 2013; Sabet et al., 2015), there was low but detectable colocalization of WT EphA2 with Rab5 in early endosomes in the absence of ligand, which was significantly increased after 15 min of ligand exposure (**Fig. 4F**). Following 60 min treatment, most WT EphA2 entered into late endosomes marked by Rab7 (**Fig. 4G**). In contrast, the LS EphA2 mutant was largely refractory to ligand-induced endocytosis, with no colocalization with Rab5 or Rab7 (**Fig. 4F, G**). The FN2 mutant had little colocalization with Rab5 in the absence of ligand (**Fig. 4F**), consistent with the lack of constitutive recycling (**Fig. 4E**). However, robust internalization and colocalization with Rab5 and Rab7 were seen with FN2 mutant upon stimulation with ligand (**Fig. 4F, G**).

Together with the biochemical analysis (**Fig. 4A)**, these results demonstrate that the core assembly mediated by the symmetric interfaces is necessary and sufficient for ligand-induced canonical signaling and endocytosis of EphA2, while auxiliary assembly mediated by asymmetric contacts is dispensable. However, asymmetric contact promotes noncanonical signaling, and both symmetric and asymmetric interactions are required for autorecycling.

### 5. EphA2 Oligomerization Regulates Cell Rounding and Migration

Cell-cell contact-induced Eph-ephrin interactions often result in cell repulsion, although adhesion can also take place (Pasquale, 2008). Consistent with this, ephrin stimulation *in vitro* leads to cell rounding and inhibition of cell migration (Miao et al., 2000; Miao et al., 2009; Stallaert et al., 2018). Time-lapse imaging experiments showed that treatment of HEK293 cells expressing WT-EphA2 with ephrin-A1 led to rapid cell rounding, while vector control cells that express low levels of endogenous EphA2 were minimally affected (*Supplemental movies*, and **Fig. 5A**). Cells expressing EphA2 FN2 mutant remained responsive to cell rounding upon ligand stimulation, whereas cells expression LS mutant became refractory consistent with its resistance of canonical signaling.

**Figure 5.**
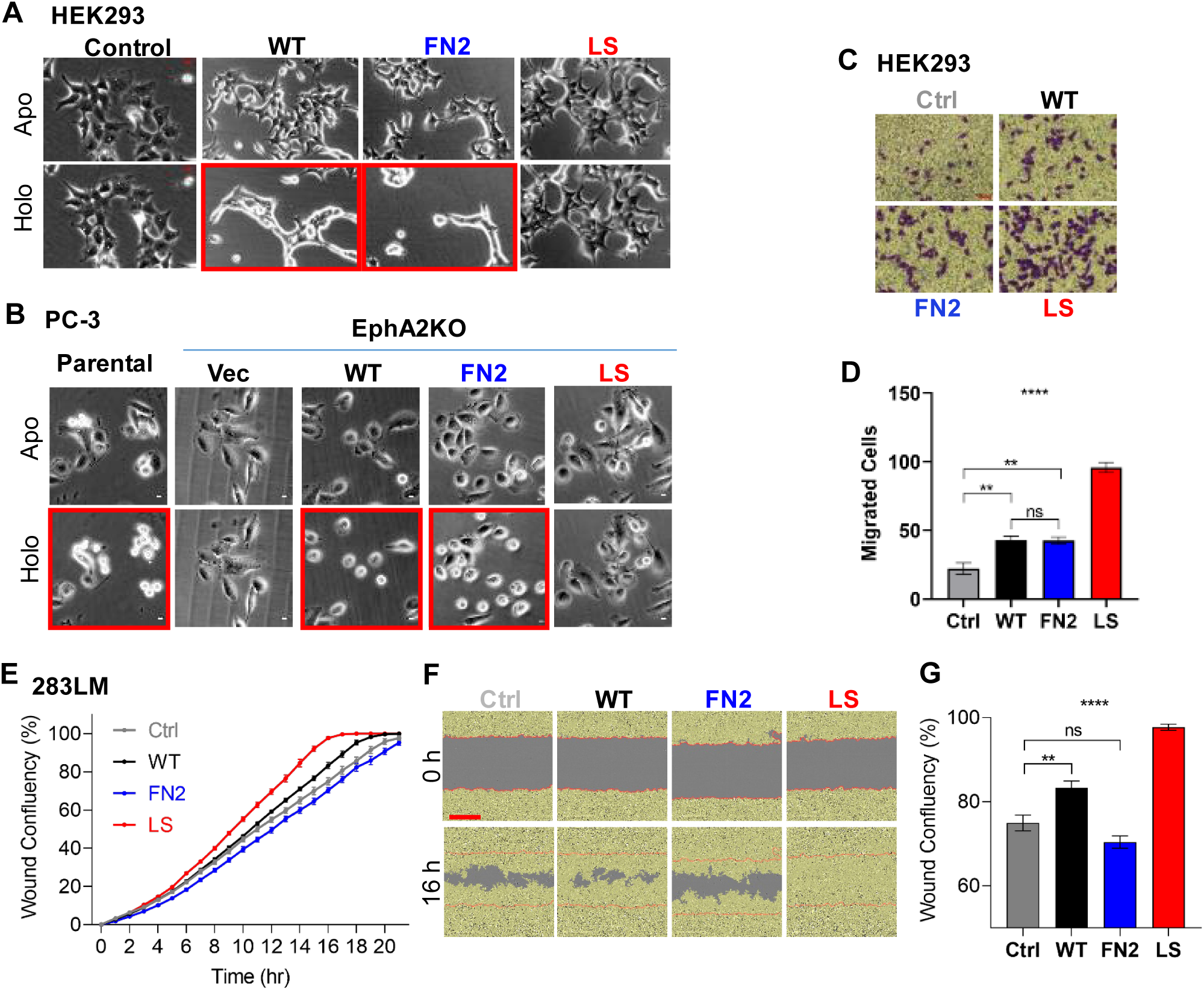
Symmetric and Asymmetric Contacts of EphA2 Modulate Cell Migration. **A)** Representative images of HEK293 cells expressing WT or mutant EphA2 upon ligand stimulation. Images are acquired at 0 and 20min after ligand stimulation. Scale bar: 5 μm. **B)** PC-3 cell with CRISPR/CAS9 knockout of EphA2 (A2KO) are reconstituted with WT or the indicated mutant EphA2. Cell images are acquired at 0 and 10min after ligand stimulation. Scale bars: 20 μm. **C,D)** Transwell cell migration of HEK293 cells expressing WT or mutant EphA2. Images are acquired 4h after seeding the cells. Scale bar: 10 μm. Number of migrated cells are counted and shown in (**D**). The error bars represent SEM. **E-G)** Two-D scratch-wound closure assay using 283LM cells expressing the indicated EphA2 constructs. Quantitative analysis is carried out on the IncuCyte metrics (**E**). Representative images of wound closure at 0 and 16 h are shown in (**F**). The yellow masks define the area covered with cells. The red lines demarcate the initial areas lacking cells. Scale bar: 400 μm. The degree of wound confluency at 16 h time point is shown in (**G**). The error bars represent SEM. The one-way ANOVA tests were performed to obtain p values (****: p < 0.0001; **: p < 0.01).

PC3 human prostate cancer cells express high levels of endogenous EphA2 and were the first cell type reported to round up in response to ligand stimulation (Miao et al., 2000). We found that PC3 cells with CRISPR/CAS9 KO of EphA2 (**Fig. S5A**) completely lost their cell rounding response. Moreover, reconstitution of the WT or FN2 mutant EphA2, but not the LS Eph mutant, restored the responsiveness (**Fig. 5B**). Similar results were obtained with SCC728 cells derived from *EphA1/EphA2* double KO mouse after re-expressing WT, FN2 or LS EphA2 (**Fig. S5B**). Together, these data demonstrate that EphA2 is a dominant mediator of ligand-induced cell rounding in PC3 cells and that the two symmetric interfaces are required for this response.

Using a 3D chemotactic cell migration assay, we found that overexpression of the LS mutant in HEK 293 cells strongly promoted chemotactic cell migration compared to the WT or FN2 mutant receptors (**Fig. 5C, D**). Next, we performed a 2D scratch wound cell migration assay using a cutaneous squamous carcinoma cell line (283LM) derived from a skin tumor induced by DMBA/TPA in an *Efna1, Efna3* and *Efna4* ligand gene triple knockout (TKO) mouse (Miao et al., 2015) (Guo et al., 2006). Since the scratch wound assay is performed with freshly confluent cells, the interactions of EphA2 with endogenous ephrin-A ligands on neighboring cells can complicate data interpretation. Using cells from the TKO mice significantly mitigate this concern. Endogenous EphA2 was knocked out from 283LM cells by CRISPR/CAS9, and WT and mutant exogenous EphA2 were reintroduced (**Fig. S5C**). We found that overexpression of WT EphA2 promoted basal 283LM cell migration. Expression of the FN2 mutant reduced the 2D cell migration (**Fig. 5E-G**), whereas the LS mutant enhanced it (**Fig. 5E-G**). Since only the asymmetric LBD-FN2 interface remains in the LS mutant (**Fig. 3F**), the stimulatory effects on both 2D and 3D migration show that EphA2 receptors in the auxiliary flanks promote cell motility, possibly by keeping the ICDs spatially apart as revealed by homoFRET (**Fig. S2E,F**) and thereby increasing noncanonical signaling.

### 6. Loss of the Symmetric Eph Core Contacts Promotes Diffuse Infiltrative Invasion *in vivo* and Shortens Host Survival in a Syngeneic Murine Glioma Model

EphA2 is a known oncogenic driver in glioma, in part by promoting diffuse infiltrative invasion(Binda et al., 2012; Miao et al., 2015), a major cause of poor prognosis of the disease. A TCGA database search reveals that EphA2 is poorly expressed in the normal brain but is markedly upregulated in GBM, particularly in the Mesenchymal and Classical molecular subtypes (**Fig. S6A)**, and the overexpression has been correlated with poor overall survival (Binda et al., 2012). Using the murine GBM cell line 1816, which lacks expression of *Nf1* and *Tp53* (Gursel et al., 2011; Pan et al., 2017; Reilly et al., 2000) and shares molecular signature of human Mesenchymal GBM, we examined the roles of EphA2 oligomerization in gliomagenesis following intracranial implantation into syngeneic C57Bl/6 mice. The choice of the immune competent syngeneic model over the conventional human xenograft in the immune deficient model was prompted by the recently reported role of EphA2 in repulsing CD8 cytotoxic T cells from the tumor mass and thus promoting tumor development (Markosyan et al., 2019).

As shown in **Fig. 6A**, 1816 cells overexpressing WT EphA2 showed reduced survival compared to parental cells, similar to previous reports (Binda et al., 2012; Miao et al., 2015). Notably, expression of the FN2 mutant EphA2 showed improved survival relative to WT. This observation correlates with the inhibitory effects of the FN2 mutation on cell migration *in vitro* (**Fig. 5E-G**). In contrast, the LS mutant that enhanced non-canonical signaling and stimulated cell migration *in vitro,* caused worse survival compared to the WT and FN2 mutant EphA2 (**Fig. 6 A, B**).

**Figure 6.**
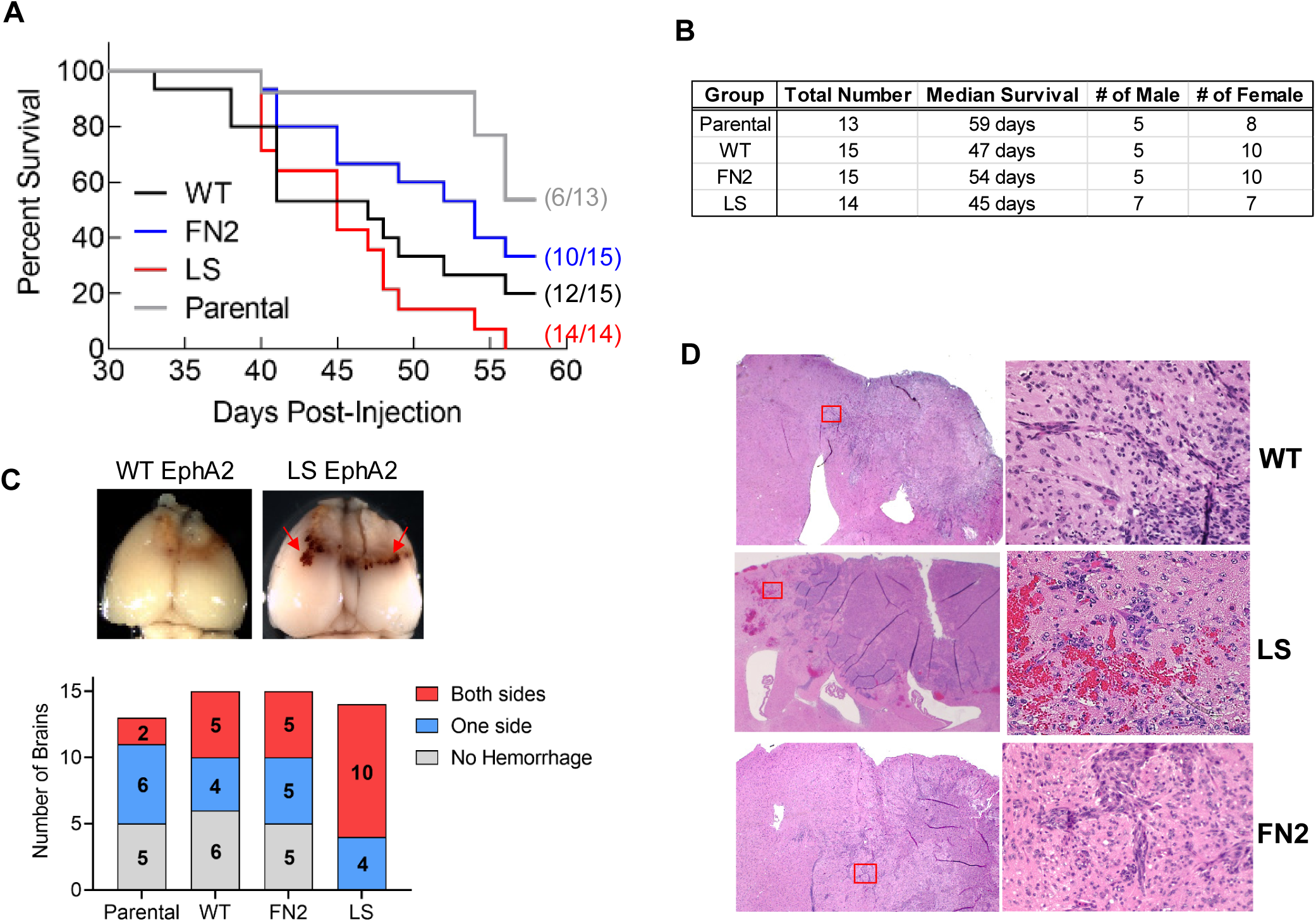
The Effects of Symmetric and Asymmetric Contacts of EphA2 *in vivo*. **A)** Kaplan-Meier survival curve of syngeneic C57Bl/6 mice injected intracranially with 1816 cells expressing the indicated WT or the indicated mutant EphA2. **B)** Number, gender and median survival of mice. **C)** Top: Representative whole mount brain images. Arrows point to regions of hemorrhage. A full collection of whole mount images is shown in **Fig. S6B**. Bottom: Numbers of mice with hemorrhage and biospheric tumor spreading from gross examination of the whole brain. **E**) Histology evaluation of mouse brains. Left panels: low power view of the brain; Right side: inset from the left.

Whole brain examination revealed that mice receiving cells expressing the LS EphA2 mutant showed a high frequency of tumor cells invading across the midline to the other hemisphere of the brain (10/14), which was accompanied by uniform presence of hemorrhage (**Fig. 6C**, and **Fig. S6B**). Invasion across the midline occurred at much lower frequencies for the tumors with WT and FN2-mutated EphA2, and hemorrhage was milder and presented less often. Histological analysis confirmed extensive spreading of the tumor cells expressing the LS EphA2 mutant to the other hemisphere with hemorrhage at the periphery of the tumor mass, whereas the tumors containing the WT or FN2 mutant EphA2 were often restricted to one side of the brain (the site of tumor cell implantation) with less prominent hemorrhage (**Fig. 6D**). These data suggest that the multimeric assembly of EphA2 receptor regulates malignant invasive behaviors *in vivo*. Disrupting the core of the Eph oligomeric assemblies promotes invasion and reduces host survival *in vivo* by ablating canonical and promoting noncanonical signaling, while disrupting the auxiliary assemblies improves the host survival by attenuating noncanonical signaling.

## Discussion

The results from the biophysical, cellular and biochemical studies reported here support a model for EphA2 assembly depicted in **Fig. 7**. In the absence of ligand binding, EphA2 apo receptors are assembled into multimers via the symmetric (LBD-LBD and Sushi-Sushi) and asymmetric (LBD-FN2) interfaces (**Fig. 7A**). Engagement via symmetric interfaces forms the core of the Eph multimer, and the interaction via the asymmetric interface extends multimerization through auxiliary assembly on the flanks. While two receptors on each side are shown in the model, the number of auxiliary receptors could vary, depending on other competing interactions, such as Ephrin-Eph in cis interactions (Kao and Kania, 2011; Seiradake et al., 2010) (Falivelli et al., 2013) (Carvalho et al., 2006). This assembly keeps kinase domains in ICD apart to facilitate the ligand-independent noncanonical signaling through Akt-, RSK1- and PKA-mediated phosphorylation at S897. Upon ligand stimulation, the asymmetric FN2-LBD interactions are displaced by high-affinity ligand-receptor (ephrin-LBD) interactions (**Fig. 7B**). Seiradake et al. reported a 71°rotation of the FN2 domain relative to the rest of the Eph ECD (FN1-CRD-LBD, which is structurally rigid) upon ligand-binding (Seiradake et al., 2010). This reorientation at the hinge-like FN1-FN2 linker would facilitate the recruitment of additional receptors into the EphA2-ephrin clusters (**Fig. 7B**). The conformational changes in ECD are propagated to ICD to induce rearrangement of the kinase domains into close proximity for transphosphorylation on tyrosines. These clusters then undergo lateral condensation into large EphA2-ephrin higher-order clusters (**Fig. 7C**) to achieve ligand-dependent canonical signaling, while the noncanonical signaling through phosphorylation of S897 is attenuated (**Fig. 7D**).

**Figure 7.**
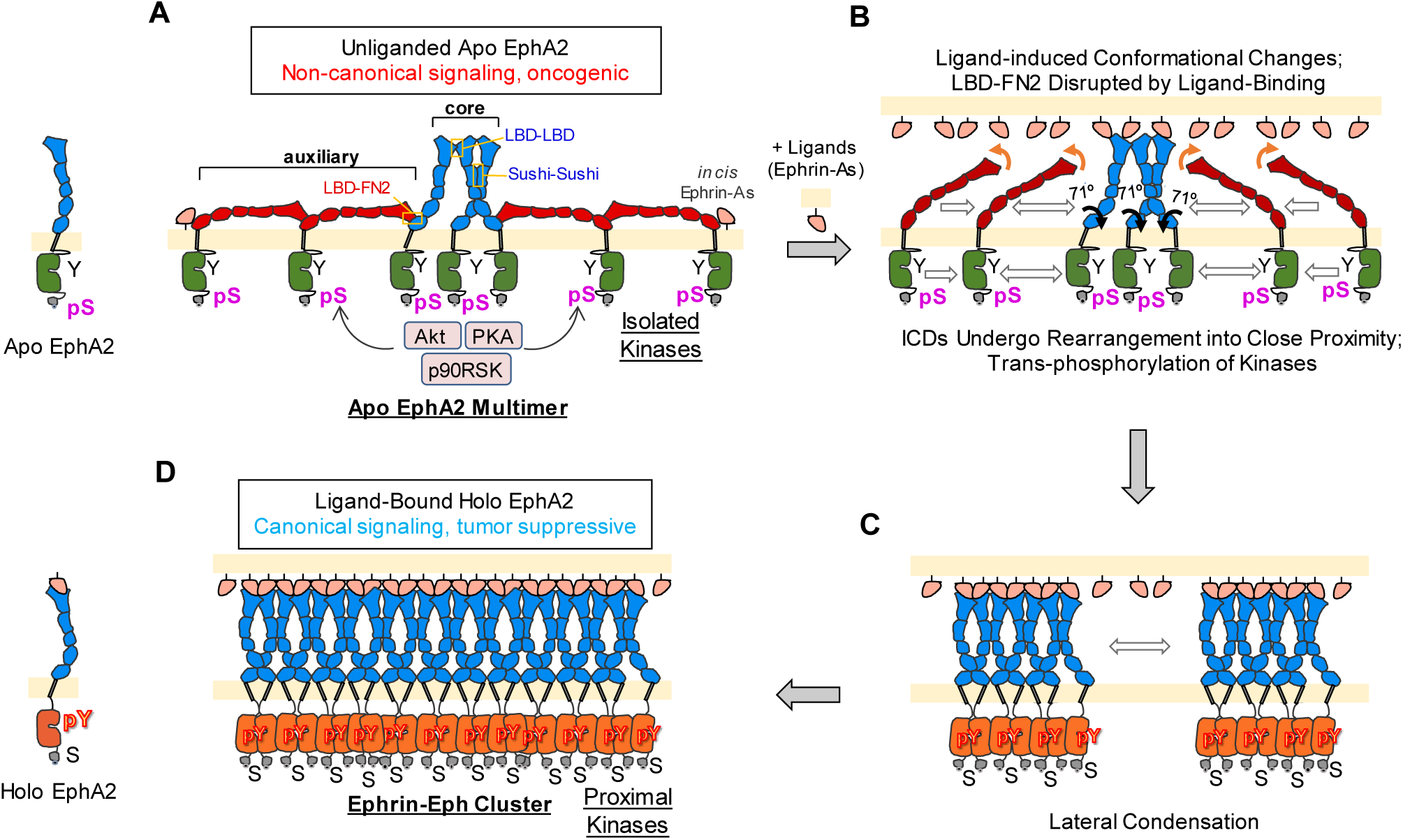
Schematic depiction of the molecular assembly of EphA2 on cell surface. See text for details.

The intricate assembly of ligand-free EphA2 apo receptor multimers reported here represents a new paradigm for RTK cell-surface organization (**Fig. 7A**). The multimeric Eph assembly contrasts with other RTKs that exist either as monomers or dimers in the absence of ligand. For example, previous investigations using the same PIE-FCCS technology shows that the EGFR apo receptor is present predominantly as a monomer on cell surface(Endres et al., 2013). Dimerization of apo receptors has been observed in several RTKs through different molecular mechanisms. For example, the transmembrane region mediates the ligand-independent dimerization of DDR1 and DDR2 (Carafoli and Hohenester, 2013), whereas the insulin receptor and the closely related IGF1 receptor are both expressed on the cell surface as preexisting disulfide linked dimers (Schlessinger, 2014). The EphA2 apo receptor is the first found to assemble into multimers. Moreover, the assembly serves essential functions in regulating auto-recycling, endocytosis, as well as canonical and noncanonical signaling.

The interactions between EphA2 apo receptors have been previously examined by different FRET assays, leading to reports of either monomers or dimers depending on the system and method used (Sabet et al., 2015; Singh et al., 2015). FRET measurements depend on the proximity of the fluorescent tags on EphA2 and the typical Förster radius for fluorescent proteins is around 60 Å (Bajar et al., 2016). However, the length of rigid ECD of EphA2 is around 146 Å (Himanen et al., 2010), well beyond the Förster radius. Since the EphA2 receptors in the auxiliary are connected by the head-to-tail (LBD-FN2) ECD interactions (**Fig. 7A**, auxiliary), rendering them “invisible” to FRET assays. Unlike FRET, PIE-FCCS measures the co-diffusion of tagged receptors to quantify their oligomerization state and is thus compatible with more spatially distant fluorescent tags within the same molecular assembly, such as those in the auxiliary arms (**Fig. 7**). The changes in oligomerization are also corroborated by a changes of the mobility of the EphA2 assemblies, which is measured directly with PIE-FCCS. In addition to these diffusion-based readouts, PIE-FCCS measures the fluorescence lifetime, providing information on the Eph spatial arrangement within the oligomers. With each of these interconnected pieces of information (summarized in Table S1), PIE-FCCS provides a comprehensive characterization of the contribution of the ECD domains to the functional EphA2 assembly in live cell membranes.

There are 14 Eph receptors in the mammalian genome. Whether the other 13 Ephs utilize similar spatial organization is an open question. However, existing evidence suggests otherwise, at least for EphA4. Crystal structures of the EphA4 ECD showed a more rigid FN1-FN2 linker, as well as mostly dimeric receptor assemblies in the presence of ligand, which is distinct from the large array-like lattice seen for EphA2/ephrin-A5(Seiradake et al., 2013; Xu et al., 2013). Indeed, our preliminary characterization of the EphA4 apo receptor using PIE-FCCS yield *f_c_* values that are consistent with dimeric assemblies that transition into smaller multimers upon ligand binding (data not shown).

Our results indicate that the ligand-induced higher-order EphA2 clustering is primarily driven by the receptor ectodomain. Since the initial discoveries of ephrin ligands, it was generally believed that ephrins must be presented as multimers to drive Eph receptor activation. In cell-based assays, this is often achieved by super-clustering of ephrin-Fc with anti-Fc antibodies (Davis et al., 1994). While this notion has been proven true for EphB receptor activation (by ephrin-B-Fc ligands) in subsequent studies (Holland et al., 1997) (Huynh-Do et al., 1999), EphA2 can be efficiently activated by dimeric ephrin-A1-Fc (Miao et al., 2000) or even monomeric ephrin-A1 (Wykosky et al., 2008). Using PIE-FCCS, we find that eprhin-A1 is a monomer, suggesting monomeric ephrin-A1 is sufficient to activate EphA2 upon cell-cell contact under physiological conditions. On the receptor side, the fact that mutating all three Eph-Eph interfaces renders EphA2 into a monomer, irrespective of ligand engagement, demonstrates that multimerization of the apo receptor and higher order clustering of ligand-bound holo receptors are primarily mediated by homotypic EphA2 ectodomain interactions. The ectodomain swapping between EphA2 and EphA4 supports this notion, as the EphA2_ECD_/EphA4_ICD_ chimeric receptor forms large clusters on the cell surface, much like EphA2, while the EphA4_ECD_/EphA2_ICD_ chimeric receptor behaves like EphA4 forming smaller clusters (Seiradake et al., 2013).

Finally, the multimeric assemblies of unliganded EphA2 have pathological and therapeutic implications. EphA2 is overexpressed in many solid human tumors, which is often accompanied by simultaneous loss of ligand expression (Al-Ejeh et al., 2014; Ireton and Chen, 2005; Miao and Wang, 2009; Pasquale, 2010; Wykosky and Debinski, 2008), creating conditions that promote EphA2 apo receptor multimeric assembly and oncogenic signaling through S897 phosphorylation. Since the EphA2 ECD plays a dominant role in receptor multimerization, its ready accessibility makes it amenable to therapeutic interventions. We posit that the LBD-FN2 asymmetric EphA2 interface may also be a good target for therapeutic development. By disrupting the asymmetric Eph-Eph interactions, the pro-oncogenic non-canonical unliganded-EphA2 signaling could be attenuated, which can be exploited alone or in conjunction with other agents to suppress malignant tumors.

## Acknowledgments

We thank members of the Wang lab and the Smith lab for the technical advice and help. We thank Elena Seiradake for help with the structural insight on the LBD-LBD, FN1-FN1 interfaces. We thank the flow cytometry shared facilities of Case Comprehensive Cancer Center supported by P30CA043703 grant from NCI. This work was supported by NIH grants (R01NS096956, R01CA250067 and R01CA155676) to BW, AWS was supported by the National Science Foundation under grant number CHE-1753060 and a Research Grant from HFSP (Ref.-No: RGP0059/2019), and DN is supported by NIH grant R01HL134570.

## Author Contributions

XS, AWS, and BW designed the study and drafted the manuscript. XS and AWD developed and designed the PIE-FCCS experiment and carried the measurements. SYK participated in PIE-FCCS measurements. CJH, DH did the in vivo experiment and data analysis. YG and LC carried out the fluorescence anisotropy imaging experiment. JH and DN provided structural details of the LBD-FN2 interface and designed the mutations. XS, RL and PT made the EphA2 mutation constructs. XS, CC and JZ generated and tested mutant EphA2 expression cell lines. SYK, YG, LC, DH, JH, KSA and MB provided manuscript edit, VV mined TCGA database and provided analysis. BW oversaw the study and manuscript preparation. All authors contributed to the final manuscript.

