## Supplemental Figures for "Cell Surface Multimeric Assemblies Regulate Canonical and Noncanonical EphA2 Receptor Tyrosine Kinase Signaling"

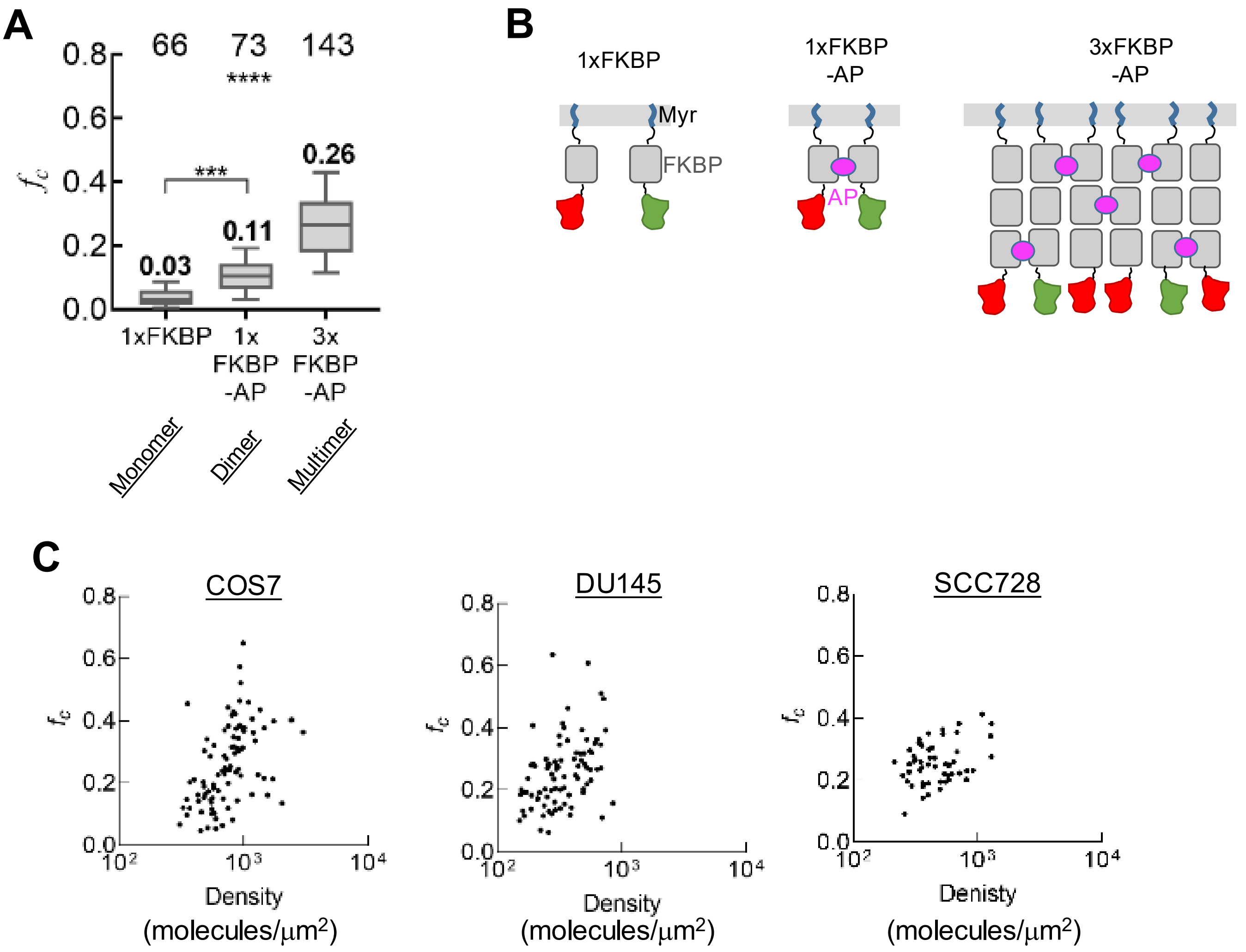


**Figure S1.** FKBP-based calibration system of cell surface oligomerization status and cell surface densities of fluorescent protein-tagged EphA2 receptors

**A)** Cross-correlation values ($f_c$) of the FKBP control constructs reported previously.

**B)** Cartoon depiction of the FKBP control constructs.

**C)** Cross-correlation values ($f_c$) of Apo EphA2 plotted against molecular density.

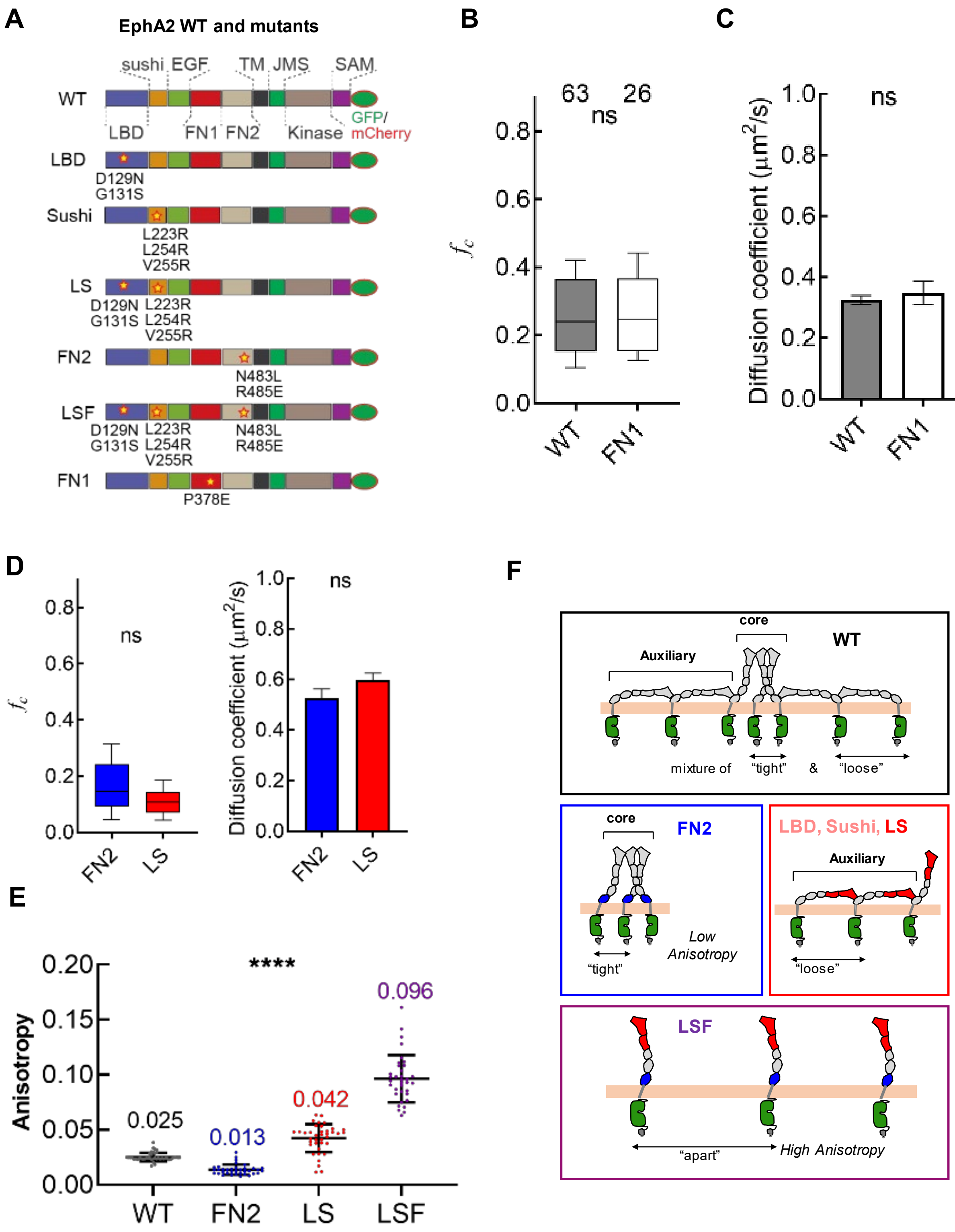


**Figure S2.**
**A)** Diagram of EphA2 constructs used in this study.
**B)** Cross-correlation values of WT EphA2 and FN1 mutant.
**C)** Diffusion coefficients of WT and FN1 mutant.
**D)** Cross-correlation values and diffusion coefficients of Apo FN2 and LS in COS 7 cells.
**E)** Anisotropy of EphA2-GFP constructs
**F)** Cartoons illustrating the different intracellular proximity caused by different extracellular contacts.

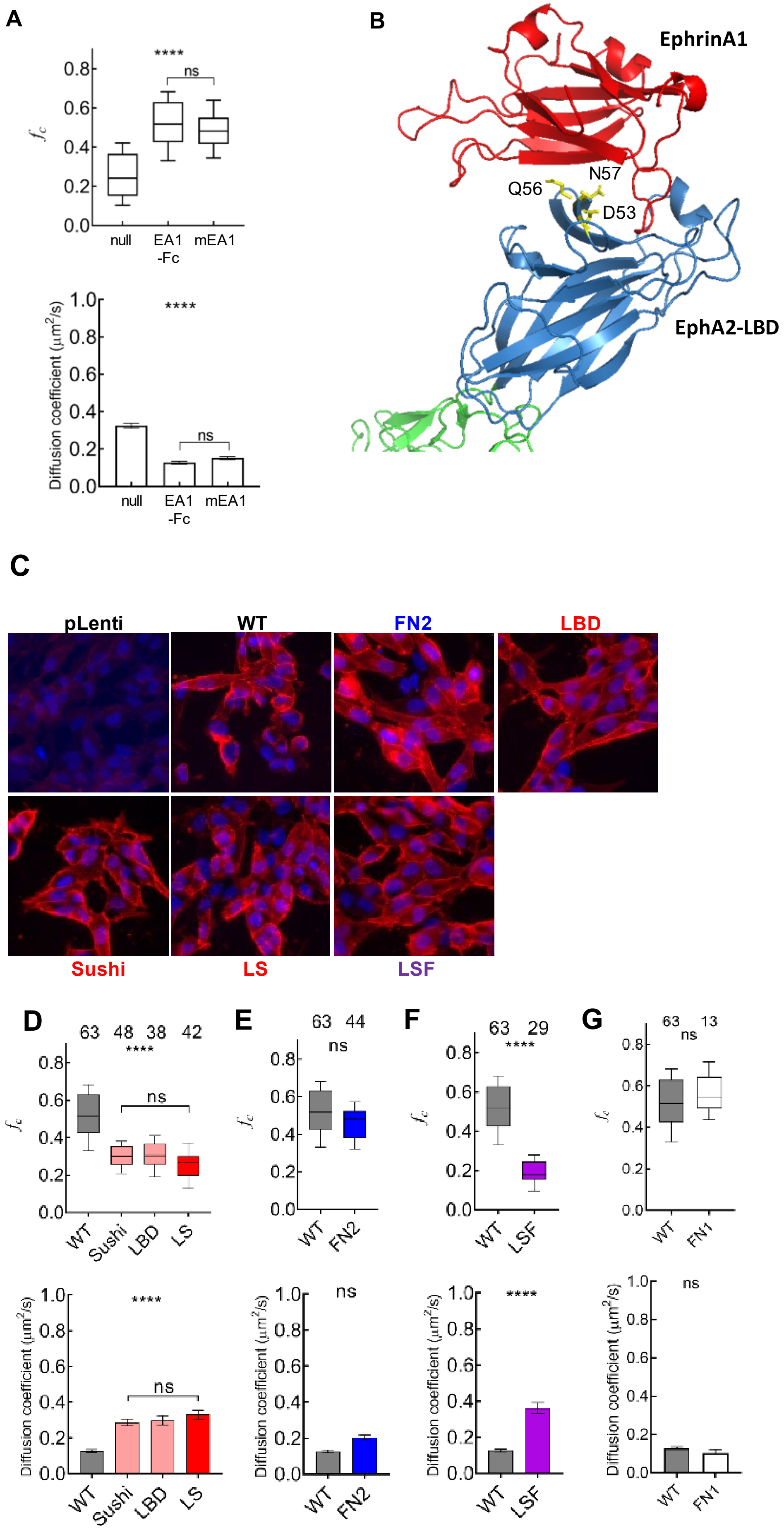
A
****
ns
0.8
0.6
0.4
0.2
0.0
fc
null
EA1 -Fc
mEA1
1.0
****
Diffusion coefficient (μm²/s)
ns
B
EphrinA1
Q56
N57
D53
EphA2-LBD
C
pLenti
WT
FN2
LBD
Sushi
LS
LSF
D
63 48 38 42
****
ns
WT
Sushi
LBD
LS
E
63 44
ns
FN2
F
63 29
****
LSF
G
63 13
ns
FN1

**Figure S3.**

**A)** Cross-correlation values (top) and diffusion coefficients (bottom) of monomeric EphrinA1 (mEA1) or dimeric Ephrin-A1-Fc WT EphA2 on Cos7 cells..

**B)** Structure of EphrinA1 binding with LBD of EphA2. The residues involved in LBD-FN2 head-tail interaction are shown in yellow sticks and labeled. This LBD-FN2 interface locates close to the ligand-binding pocket on LBD and is blocked by ligand-binding.

**C)** WT and mutant EphA2 bind to Ephrin-A1-Fc ligand equally well on GSC827 cells. Cell were incubated with 10 nM Ephrinin-A1-Fc fro 30 minutes on ice. Bound Ephrin-A1-Fc was visualized with red fluorescent anti-Fc antibodies. Scale bars: 20mm.

**D)** Cross-correlation values (top) and diffusion coefficients (bottom) of ephrinA1-Fc (EA1-Fc) stimulated EphA2 constructs (Sushi, LBD and LS) with disruption at symmetric contact.

**E)** Cross-correlation values (top) and diffusion coefficients (bottom) of EA1-Fc stimulated EphA2 construct (FN2) with disruption at asymmetric contact.

**F)** Cross-correlation values (top) and diffusion coefficients (bottom) of EA1-Fc EphA2 construct (LSF) with disruption at both symmetric and asymmetric contacts. The cross-correlation values of EA1-Fc stimulated LSF are close to dimer control, indicating that EA1-Fc stimulation causes bias in oligomerization states.

**G)** Cross-correlation values (top) and diffusion coefficients (bottom) of EA1-Fc stimulated (Holo) FN1.

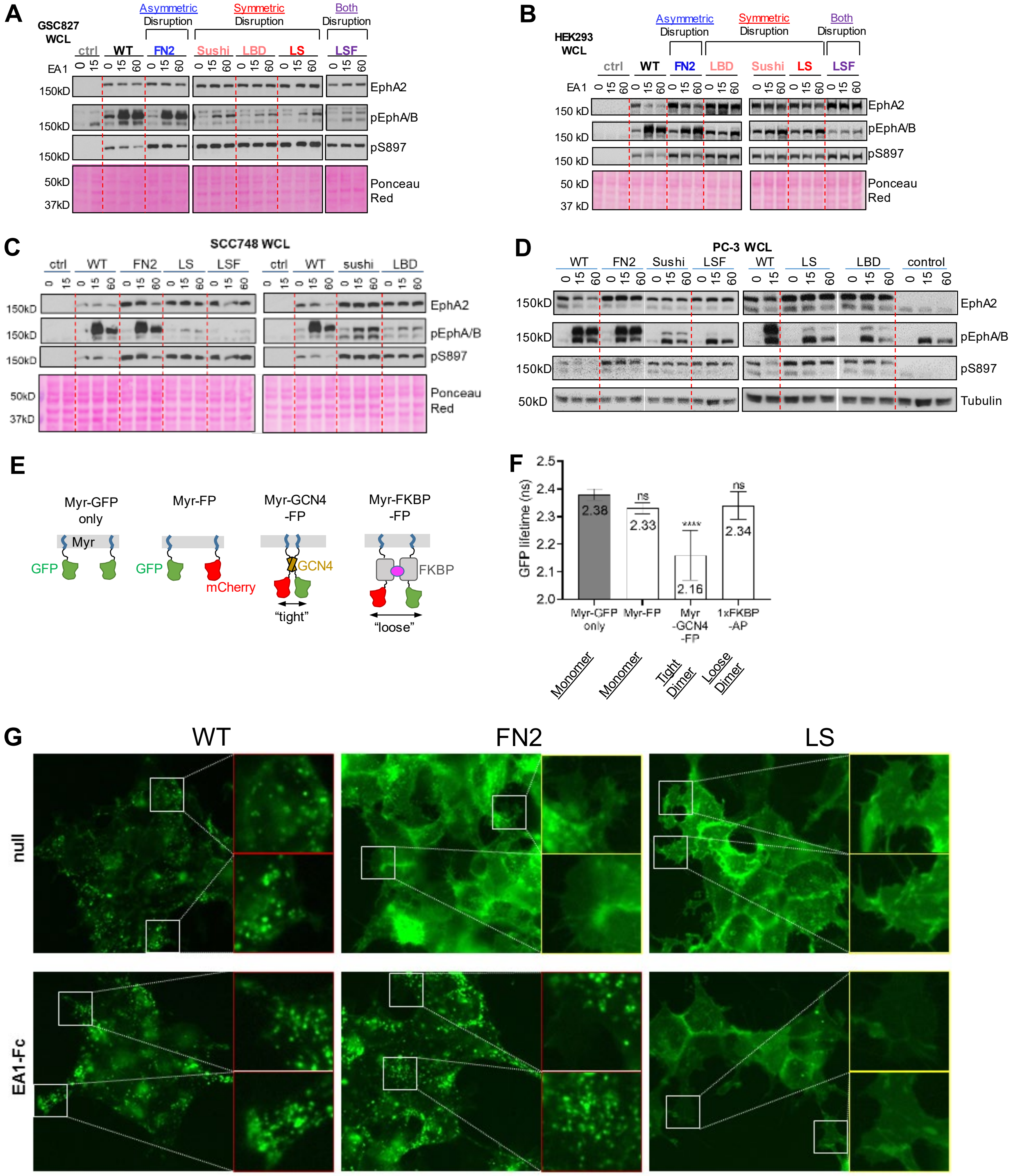


**Figure S4.**

**A, B, C, D)** EphA2 constructs are expressed in GSC827 **(A)**, HEK293 **(B)**, SCC748 **(C)** and PC-3 **(D)** cells are stimulated with 3 mg/mL ephrinA1-Fc for 15 min and 60 min, and lysed. Whole-cell lysates are subjected to immunoblot with the indicated antibodies.

**E)** Fluorescence lifetime of GFP on different control constructs. Only the tight dimer control Myr-GCN4-FP shows significantly shorter lifetime, indicating FRET from GFP to mCherry.

**F)** Cartoon depiction of the control constructs.

**G)** Epi-fluorescence images of HEK293 with different EphA2 localization patterns.

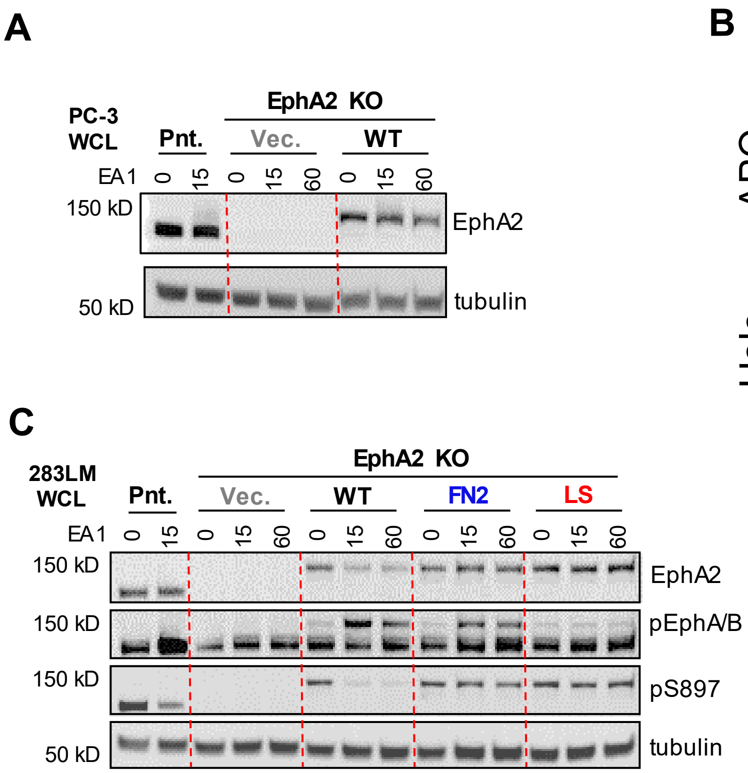


**Figure S5.**

**A)** PC-3 cells are stimulated with 3 μg/mL ephrinA1-Fc for 15 min and 60 min, and lysed. Whole-cell lysates are subjected to immunoblot with the indicated antibodies.

**B)** Example images of SCC728 cells expressing EphA2 constructs responding to ligand stimulation. Images are acquired at 0 and 20min after ligand stimulation. Cell rounding is observed with WT and FN2 expressing cells, but not with LS expressing cells. Scale bar: 5 μm.

**C)** EphA2 constructs are expressed in 283LM cells are stimulated with 3 μg/mL ephrinA1-Fc for 15 min and 60 min, and lysed. Whole-cell lysates are subjected to immunoblot with the indicated antibodies.

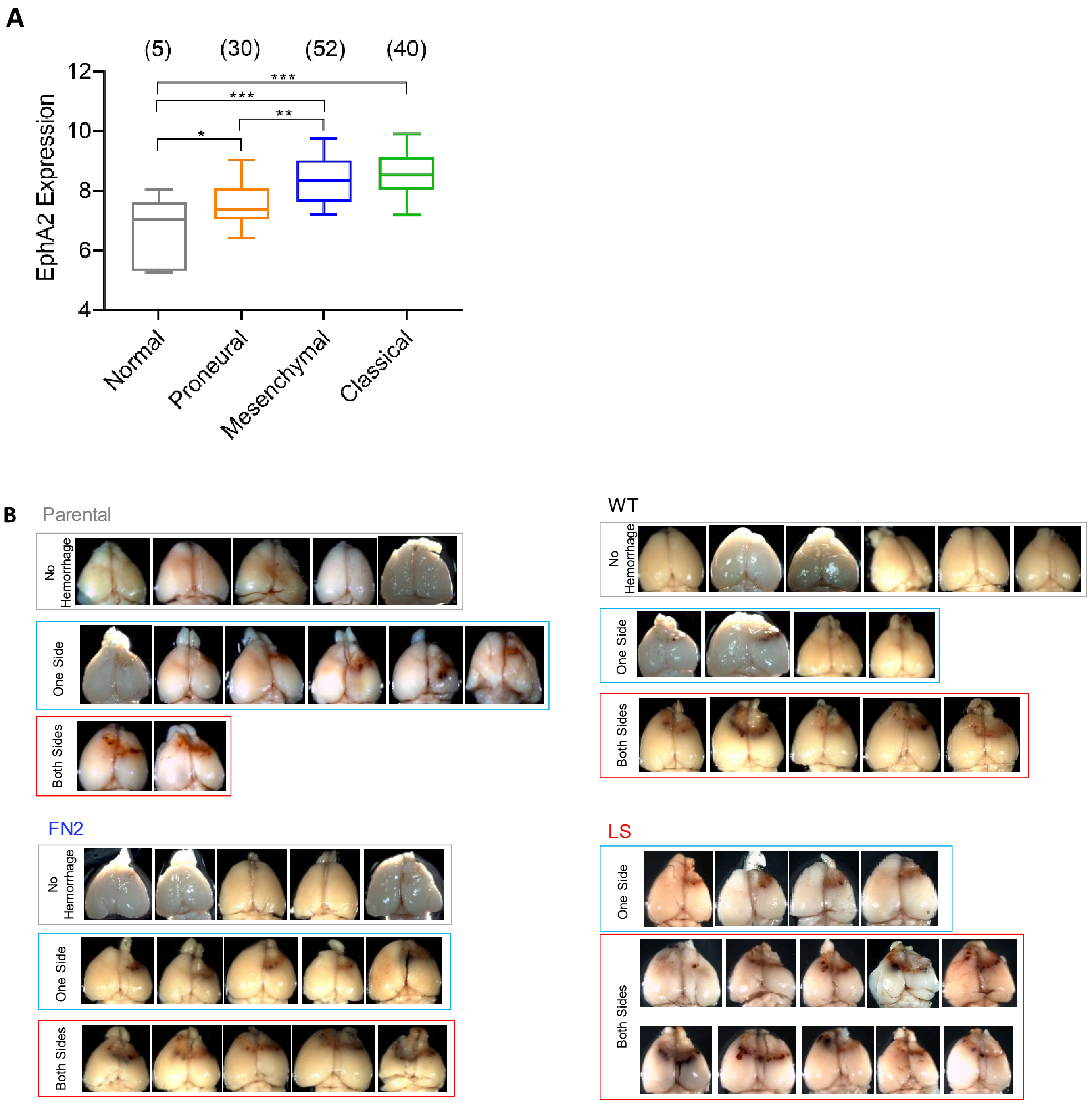


**Figure S6.**

A) Expression levels of EphA2 in different types of brain tumor.

B) Images of all the brain samples from the gliomagenesis experiment. The samples are divided into groups according to the patterns of the hemorrhage. The brains injected with LS expressing cells show most severe hemorrhages.

Supplemental Movies

Movie S1. Time lapse images of HEK293 cells expressing EphA2 constructs in response to EA1 stimulation. Note WT and FN2 expressing cells undergo significant rounding, while LS expressing cells do not.
