## Supplemental Table 1 for "Cell Surface Multimeric Assemblies Regulate Canonical and Noncanonical EphA2 Receptor Tyrosine Kinase Signaling"

**Table S1.** Summary of cross correlation value, diffusion coefficient, GFP life time and anisotropy of WT and each mutant EphA2 in the absence of presence of monomeric or dimeric ephrin-A1 ligand.

| EphA2 | Unliganded (APO) | mEphrinA1 (Holo) | EphrinA1-Fc (Holo) | Cross-correlation value *($f_c$)* | Diffusion coefficient ($\mu m^2/s$) | GFP lifetime (ns) | Anisotropy |
|---|---|---|---|---|---|---|---|
| WT | + | | | 0.24 | 0.33 | 2.13 | 0.025 |
| WT | | + | | 0.48 | 0.15 | 2.05 | |
| WT | | | + | 0.52 | 0.13 | 2.03 | |
| FN2 | + | | | 0.15 | 0.53 | 2.17 | 0.014 |
| FN2 | | + | | 0.37 | 0.24 | 2.09 | |
| FN2 | | | + | 0.48 | 0.2 | 2.07 | |
| LBD | + | | | 0.15 | 0.54 | 2.15 | |
| LBD | | + | | 0.14 | 0.62 | 2.15 | |
| LBD | | | + | 0.3 | 0.3 | 2.12 | |
| Sushi | + | | | 0.12 | 0.59 | 2.18 | |
| Sushi | | + | | 0.25 | 0.43 | 2.18 | |
| Sushi | | | + | 0.3 | 0.29 | 2.17 | |
| LS | + | | | 0.11 | 0.6 | 2.14 | 0.042 |
| LS | | + | | 0.06 | 0.74 | 2.15 | |
| LS | | | + | 0.27 | 0.33 | 2.11 | |
| LSF | + | | | 0.03 | 0.82 | 2.16 | 0.096 |
| LSF | | + | | 0.03 | 0.62 | 2.19 | |
| LSF | | | + | 0.18 | 0.36 | 2.18 | |
| FN1 | + | | | 0.25 | 0.35 | 2.17 | |
| FN1 | | + | | 0.5 | 0.15 | 2.02 | |
| FN1 | | | + | 0.55 | 0.1 | 2.03 | |
